# The binding landscape of Ras/effector interactions provides insights on Ras switching and oncogenesis

**DOI:** 10.64898/2026.11.04.764156

**Authors:** Atilio Tomazini, Sagara Gurusinghe, Sergei Bogomolov, Julia M. Shifman

## Abstract

Protein-protein interactions (PPIs) regulate essential cellular processes, yet the evolutionary principles that shape their specificity, adaptability, and dysregulation in disease remain poorly understood. Here, we investigate the evolution of a signaling PPI between the small GTPase Ras and its effector Nore1A, a tumor suppressor involved in Ras-mediated signaling. Ras acts as a molecular switch, engaging effectors in its GTP-bound active state and releasing them after GTP hydrolysis, whereas oncogenic mutations such as G12D lock Ras in the active state, promoting aberrant signaling in cancer. Using yeast surface display, deep mutational scanning, and data normalization, we systematically mapped the Ras/Nore1A binding landscape under different physiological and oncogenic states. We generated and screened ∼33,000 Nore1A single and double mutants against RasWT and RasG12D in both GTP-and GDP-bound states. Our data revealed that most Nore1A mutations produce small-to-moderate decrease in Ras affinity, while a small number of substitutions enhance affinity. Unexpectedly, GTP hydrolysis causes minimal remodeling of the interaction landscape. Conversely, the oncogenic RasG12D mutation reshapes the binding landscape, revealing many Nore1A variants with enhanced and selective affinity for mutant Ras. Molecular dynamics (MD) simulations of Nore1A–Ras variant complexes suggest that the non-optimality of the Nore1A/RasG12D binding landscape arises, in part, from the greater intrinsic flexibility of RasG12D and the pronounced reduction in flexibility induced by affinity-enhancing Nore1A mutations. These findings demonstrate that disease-associated mutations can create new evolutionary trajectories in signaling PPIs by changing interaction dynamics and provide a framework for engineering selective protein-based inhibitors targeting oncogenic Ras variants.

**Significance:** Protein-protein interactions regulate essential cellular processes, yet the evolutionary principles governing their specificity, adaptability, and dysregulation in disease remain poorly understood. By systematically measuring the effects of thousands Nore1A mutations on binding to wild-type and oncogenic Ras, we show that most mutations modestly weaken binding, while a few strengthen it. Surprisingly, GTP hydrolysis barely remodels this landscape, whereas the G12D mutation reshapes it, revealing many Nore1A variants with enhanced and selective affinity for mutant Ras. Molecular dynamics simulations suggest that these changes arise from altered Ras flexibility and its modulation by Nore1A mutations. Our findings reveal how disease mutations can create new evolutionary trajectories in PPIs and provide principles for engineering selective protein therapeutics.

---

Protein-protein interactions (PPIs) are fundamental to nearly all cellular processes, enabling the formation of dynamic complexes that regulate signalling, structural organization, transport, and metabolism. PPIs exhibit a wide range of binding affinities, and these affinities are often finely tuned to support their biological function. High-affinity interactions are essential when proteins must form stable, long-lived complexes—for example, enzyme–inhibitor pairs or toxin–antitoxin systems that rely on tight binding to ensure precise regulation (1–4). In contrast, many signaling PPIs require moderate affinities so that complexes assemble and disassemble rapidly in response to specific signals that switch their activity on and off (5–8). A prototypical example of a protein switch is Ras, a small GTPase that engages different effector proteins in a nucleotide-dependent manner to regulate a wide range of signaling pathways (9). Ras functions by cycling between an active, GTP-bound state (Ras-GTP), and an inactive, GDP-bound state (Ras-GDP) (10, 11). Ras adopts a conserved GTPase fold, featuring a centrally positioned nucleotide-binding pocket that coordinates binding of either GTP or GDP (Fig. 1A).

**Fig. 1.**
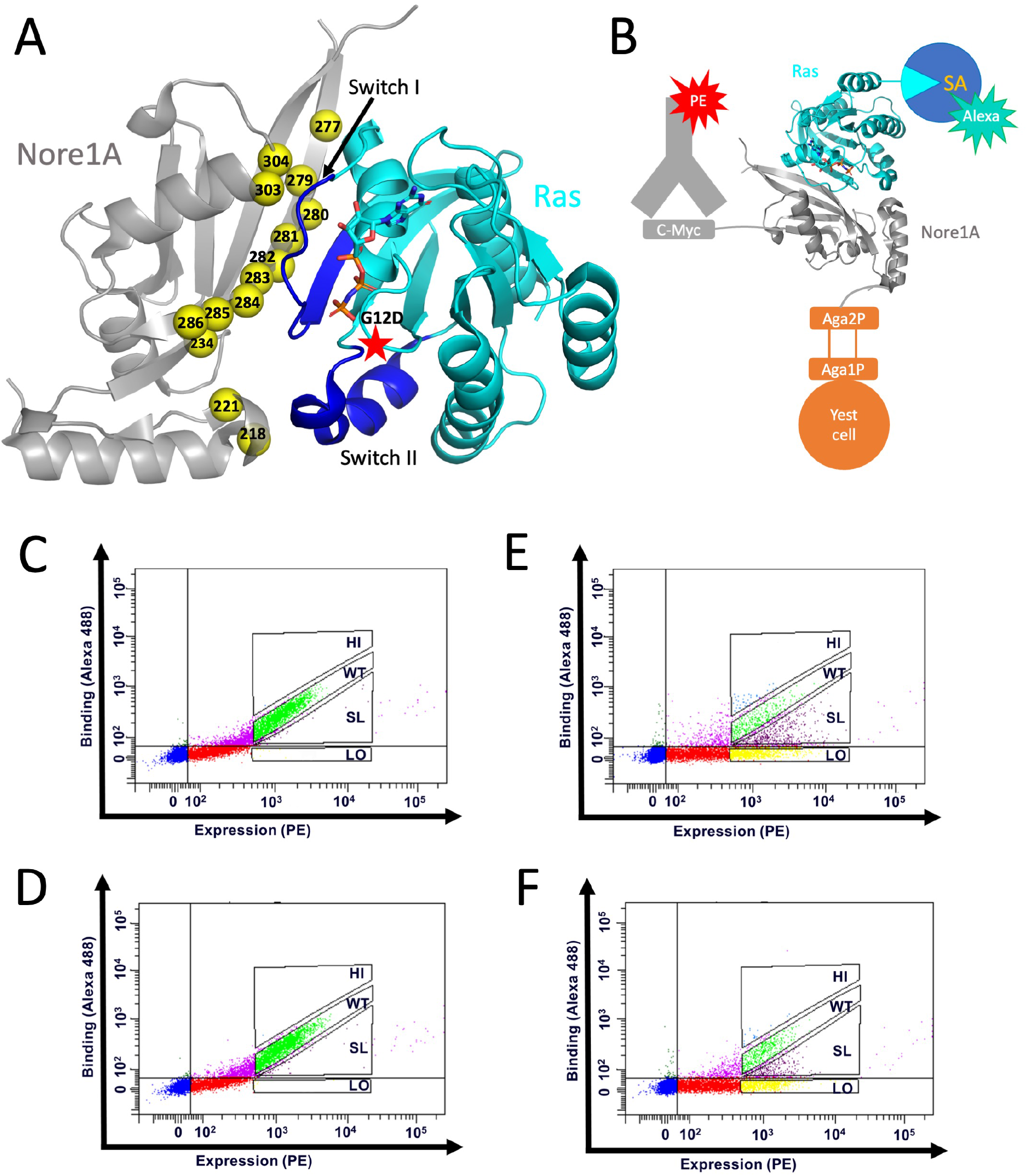
(A) Structure of the Nore1A-Ras-GTP complex (PDB ID 3DDC) showing Ras in cyan and Nore1A in grey. The nucleotide is shown as sticks, position of the G12D mutation – as a red star, and the two switches are colored blue. Nore1A positions that have been randomized in the library are shown as yellow spheres. (B) Schematic diagram showing the YSD setup used in the study. Nore1A variants are expressed as a C-terminal fusion to the Aga2P protein on the yeast surface. c-Myc tag is used to monitor Nore1A variant expression using a PE-conjugated anti-c-Myc antibody. Ras binding to Nore1A variants is monitored using Alexa488-conjugated streptavidin (SA) that binds to biotinylated Ras. (C)-(F) FACS data showing binding of Nore1A WT or Nore1A library to Ras. (C) Nore1A WT binding to RasWT-GTP (3 µM). (D) Nore1A WT binding to RasWT-GDP (15 µM). (E) Nore1A library binding to RasWT-GTP (3µM). (F) Nore1A library binding to RasWT-GDP (15µM). Four sorting gates are depicted: HI (with affinity higher than WT), WT (with wild-type like affinity), SL (with slightly lower than wild-type affinity), LO (with much lower than wild-type affinity).

Ras switching is mediated by nucleotide-dependent structural changes in two flexible regions known as switch I (residues 30–38) and switch II (residues 60–76)(Fig. 1A). In the GTP-bound state, the *γ*-phosphate of GTP stabilizes switch I and II through hydrogen bonding and electrostatic interactions, creating a high-affinity effector-binding conformation. Upon GTP hydrolysis, loss of this phosphate abolishes these interactions, allowing the Ras switches to sample a number of more open conformations. As a result, Ras-GDP must pay a large entropic penalty to adopt the effector-binding conformation, explaining its markedly reduced affinity for effectors and highlighting a central feature of Ras molecular switching (9, 12). Several point mutations in Ras, clustered around the nucleotide-binding pocket most frequently at positions 11, 12, and 61, impair the protein’s intrinsic and GAP-stimulated GTP hydrolysis, preventing efficient conversion of GTP to GDP (13). Mutations at these residues disrupt key structural elements required for Ras catalysis, effectively “locking” Ras in the GTP-bound state. As a result, Ras remains constitutively active, continuously recruiting effectors and driving uncontrolled downstream signaling that promotes cell proliferation and survival and eventually resulting in cancer (14).

Ras effectors mediate diverse cellular outcomes down-stream of Ras signaling. Among the best-characterized effectors are Raf protein kinases, which activate cell pro-liferation and differentiation(15); Ral guanine nucleotide dissociation stimulator (RalGDS), which promotes cell transformation and cytoskeletal reorganization (16, 17), and lipid kinases PI3Ks, which regulate apoptosis inhibition, cell survival, growth, and migration (18). While most Ras effectors, promote proliferative and pro-survival signaling, another Ras effector, Nore1A (also called RASSF5), drives Ras outputs in the opposite direction by activating pathways that induce senescence and cell death (19–24). This unique function makes Nore1A a compelling model for investigating the evolutionary determinants of Ras recognition and its ability to discriminate between nucleotide-bound states and oncogenic variants.

Structurally, Ras effectors are multidomain proteins that contain Ras-association (RA) or Ras-binding domain (RBD), which adopts a ubiquitin fold (25) (collectively referred to as RBD in this manuscript). All structurally characterized effectors bind Ras through a conserved interface involving an antiparallel intermolecular β-sheet formed between β2 of the RBD and β2 of Ras, together with contacts to the Ras switch I region. (Fig. 1A)(26, 27). The interactions between Ras and canonical RBDs are predominantly polar, involving multiple salt bridges and hydrogen bonds across the binding surface (28, 29). Nore1A is one of the few effectors that contacts both switch I and switch II, resulting in an expanded interaction interface and prolonged association time compared with other Ras effectors (Fig. 1A) (26).

While the structural features of Ras–effector recognition have been extensively characterized, the contribution of effector sequence variation to Ras recognition and specificity remains poorly understood. Several deep mutational studies have examined how different mutations in Ras affect Ras affinity for its effectors and modulate downstream signaling output (30–32). These studies revealed the basis for the high sequence conservation of Ras and demonstrated how oncogenic mutations reshape its fitness landscape by perturbing the allosteric network connecting Ras residues. To complement this evolutionary perspective on Ras–effector interactions and to better understand how Ras effectors recognize and discriminate between distinct Ras states, we systematically introduced mutations into the Ras effector Nore1A. Using the recently developed methodology (33), we constructed a quantitative binding landscape of the Nore1A–Ras interaction and determined how this landscape changes upon GTP hydrolysis and upon introduction of the G12D mutation, the most frequent Ras mutation in pancreatic and colorectal cancer. Our study shows that most mutations in Nore1A have a mildly destabilizing effect on Ras binding, and that only a limited number of mutations enhance its affinity for Ras, underscoring the high degree of evolutionary optimization of this interaction. More-over, the binding landscape remains largely unchanged upon GTP hydrolysis, whereas the oncogenic RasG12D mutation considerably reshapes it, revealing numerous affinity-enhancing mutations that preferentially target RasG12D over wild-type Ras. These findings point to substantial alterations in Ras–effector interactions upon oncogenic mutations and suggest potential strategies for designing mutation-specific Ras inhibitors.

## Results

### Nore1A library construction and binding selections

To map the binding landscape of the Ras/Nore1A interaction, we first incorporated the wild-type Nore1A (Nore1A WT) gene into the pCTCON2 vector, compatible with Yeast Surface Display (YSD) technology. Using this construct, Nore1A WT was displayed on the surface of yeast cells with a C-terminal Myc tag, enabling detection of protein expression via binding of an anti-c-Myc antibody followed by a phycoerythrin (PE)-conjugated secondary antibody (Fig. 1B). Wild-type HRas (RasWT) and its G12D mutant (RasG12D) was expressed in a biotinylated form, purified, and loaded with either the non-hydrolysable GTP analogue GppNHp (hereafter referred to as GTP) or with GDP. Binding of Ras to Nore1A WT was measured by monitoring the fluorescence of Alexa Fluor 488, which was linked to biotinylated Ras through streptavidin (Fig. 1B). The assessment of binding of Nore1A WT to RasWT-GTP by Fluorescence-Activated Cell Sorting (FACS) showed a diagonal narrow distribution, demonstrating that Nore1A WT is well expressed on the surface of yeast cells, is properly folded, and binds to RasWT-GTP (Fig. 1C). Similar analyses of Nore1A WT binding to RasWT-GDP, RasG12D-GTP, RasG12D-GDP were performed (Fig. 1D and Figure S1 in SI Appendix).

We subsequently generated a comprehensive library of Nore1A variants, encompassing all single and double mutants at 14 positions that form the direct binding interface with Ras in the Nore1A–Ras complex structure (24) Fig. 1A. This library, hereafter referred to as the naïve library, comprised 266 single mutants as well as all possible pairs of these mutations, resulting in a theoretical diversity of 33,118 unique Nore1A sequences. The naïve library of Nore1A mutants was transformed into yeast and analyzed by Next-Generation Sequencing (NGS), revealing complete coverage of single mutants and 51.8% coverage of double mutants after a cutoff of 10 sequencing events was applied. The majority of detected double mutants combined mutations that were distant in sequence, reflecting a bias introduced by the library-construction method. Independent NGS sequencing of two identical Nore1A naïve library samples produced highly-correlated results, with an R value of 0.984 between the two samples (Figure S2 in SI Appendix).

In the subsequent step, the Nore1A library was displayed on the yeast surface, and its expression and binding to four Ras isoforms (RasWT-GTP, RasWT-GDP, RasG12D-GTP and RasG12D-GDP) was assayed using FACS. In each experiment, the Ras concentration was adjusted to produce a similar mean fluorescence signal for Nore1A WT while allowing the Nore1A library to span a broad range of binding signals, reflecting the diversity of Ras-binding affinities among the variants, both above and below that of WT. Next, we performed a sorting experiment, collecting yeast cells with Nore1A variants that fell into four affinity groups: higher-than-WT affinity (HI), WT-like affinity (WT), slightly lower-than-WT affinity (SL), and strongly lower-than-WT affinity (LO) (Fig. 1E and F and Figure S1B, D in SI Appendix). Using four affinity gates, rather than selecting only the highest-affinity population, improves the robustness of the analysis by minimizing errors arising from cell-sorting variability and differences in mutant expression, as demonstrated in our previous study (33). Cells from each affinity gate were subsequently expanded and analyzed by NGS, yielding approximately 3 million reads per population. For each Nore1A mutant identified in the NGS data, we calculated an enrichment value for each affinity gate, representing its enrichment in that gate relative to the naïve library (see Methods). The enrichment values for single Nore1A mutants interacting with the four Ras isoforms are summarized in Fig. 2A for high-affinity (HI) gate and in Figure S3 in SI Appendix for all gates.

**Fig. 2.**
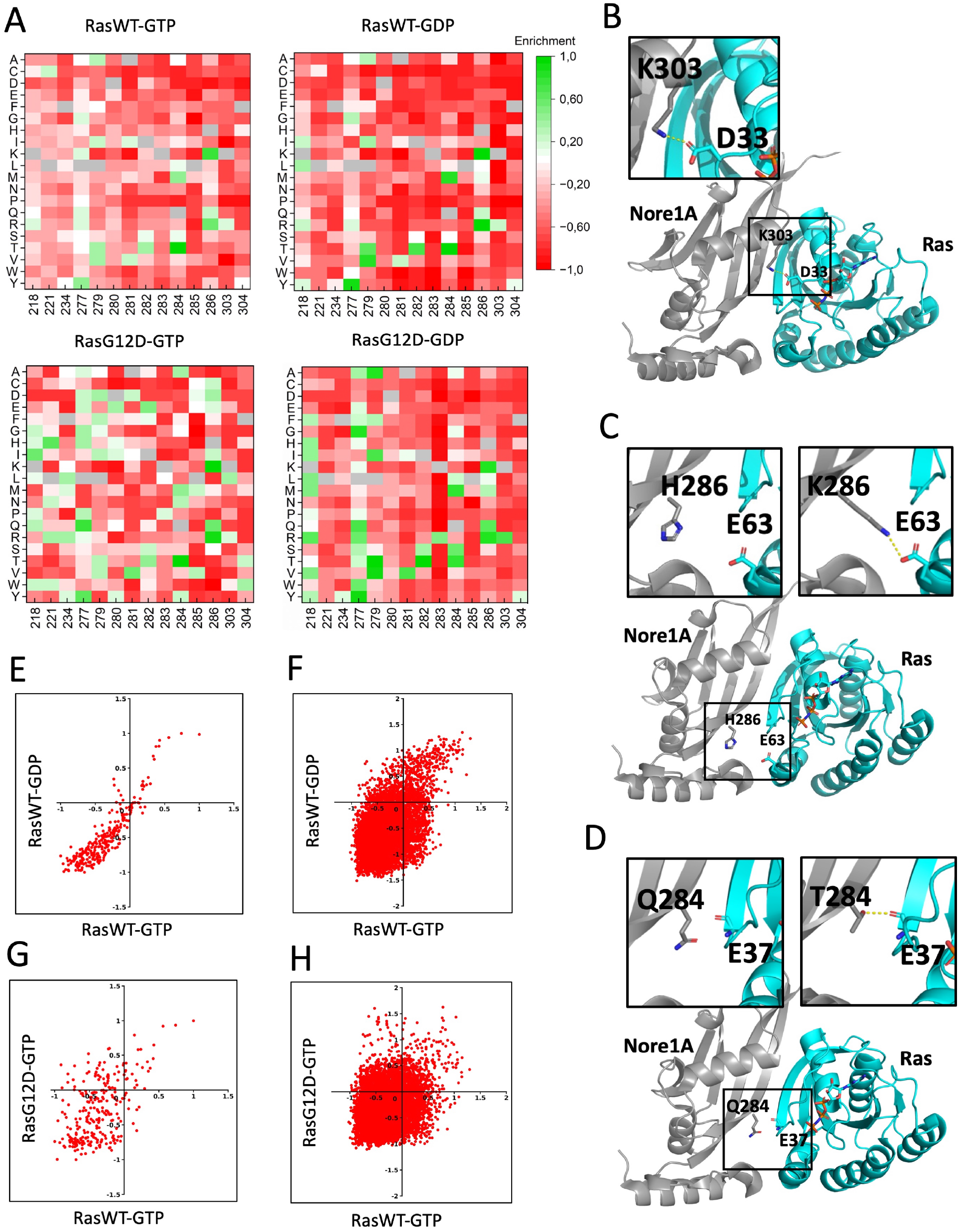
(A) Heatmaps showing normalized enrichment values in the HI gate for Nore1A single mutants binding to four Ras variants: RasWT-GTP (top left), RasWT-GDP (top right), RasG12D-GTP (bottom left), and RasG12D-GDP (bottom right). The 20 amino acids are shown on the y-axis, and the randomized Nore1A positions are shown on the x-axis. Enrichment values are color-coded from +1 (green) to 1 (red). (B) Structure of the Nore1A–RasWT-GTP complex highlighting the hotspot interaction involving K303, with a magnified view of the interaction shown above. (C) Structure of the Nore1A–RasWT-GTP complex highlighting the cold-spot residue H286. Magnified views compare the WT interface (left) with the interface following the H286K mutation (right). (D) Structure of the Nore1A–RasWT-GTP complex highlighting the affinity-enhancing Q284T mutation. Magnified views compare the WT interface (left) with the interface following the Q284T mutation (right). (E–H) Pairwise correlations between normalized enrichment values in the HI gate for different datasets: (E) RasWT-GTP versus RasWT-GDP for single mutants; (F) RasWT-GTP versus RasWT-GDP for double mutants; (G) RasWT-GTP versus RasG12D-GTP for single mutants; and (H) RasWT-GTP versus RasG12D-GTP for double mutants.

### The Nore1A–RasWT–GTP Binding Landscape Is Evolutionarily Optimized and Reflects Interface Structural Features

Our data clearly show that most single Nore1A mutations are depleted in the HI gate and enriched in the SL and LO gates (Fig. 2A top left and Figure S3 in SI Appendix). This data suggests that, given the specific geometry of the Nore1A–RasWT-GTP interface, the native Nore1A sequence is already nearly optimal for interaction with Ras and could be improved by only a limited number of mutations. Furthermore, the enrichment data closely reflect the structural features of the Nore1A–Ras interface. For instance, position 303 emerges as a clear binding hotspot in our enrichment data, with substitutions of a native lysine showing strong depletion in the HI gate (red) for all amino acids except arginine (Fig. 2A, top left). Consistent with this finding, K303 forms an intermolecular salt bridge with D33 on Ras (Fig. 2B), and arginine is the only substitution that can preserve this interaction. In agreement with our data, replacing K303 with alanine was documented to result in a large decrease in experimentally measured Nore1A affinity for Ras (24). Similarly, position 283 emerged as another hotspot in our data (Fig. 2A, top left). This position is occupied by Lys in Nore1A WT, which forms a buried salt bridge with Asp38 of Ras; disrupting this interaction results in a very substantial destabilization of the complex (24).

In contrast, position 286 emerged as a cold spot of binding (34), with the presence of three affinity-enhancing mutations in the HI gate (Fig. 2A, top left). Structural analysis shows that this position occupied by histidine does not directly interact with RasWT-GTP in WT Nore1A but could establish a new intermolecular hydrogen bond with E63 or Y64 on Ras upon substitution with Lys, Arg, or Gln (Fig. 2C). Another affinity-enhancing mutation identified by our enrichment data is Q284T. Structural modeling suggests that this substitution creates an additional hydrogen bond between Thr284 of Nore1A and the backbone CO on the opposing β-sheet of Ras, thus stabilizing the intermolecular beta-sheet (Fig. 2D).

### Nucleotide exchange does not substantially affect the Nore1A/Ras binding landscape

We next analyzed how GTP hydrolysis affects the Nore1A/RasWT binding landscape by comparing the enrichment values obtained in the RasWT-GTP and RasWT-GDP binding selections (Fig. 2A top left and right, respectively). Our data clearly show that the landscape for RasWT-GDP is very similar to that of RasWT-GTP. Indeed, enrichment values in the HI gate for RasWT-GTP and RasWT-GDP were highly correlated for single mutations (R = 0.94) (Fig. 2E). Only a few single mutations slightly deviate from the overall correlation, showing improved affinity for RasWT–GDP but not for RasWT–GTP. A similar diagonal trend is observed for double mutations, indicating that most variants have comparable effects on RasWT-GDP and RasWT-GTP binding; however, the increased scatter suggests greater variability among double-mutant variants compared to single mutants (Fig. 2F).

### G12D Alters the Nore1A-Ras Binding Landscape

Comparison of the heat maps for Nore1A interactions with RasWT-GTP and RasG12D-GTP reveals several differences in the effects of mutations on binding. Most notably, a greater number of mutations are enriched in the HI gate for RasG12D-GTP than for RasWT-GTP, suggesting that the Nore1A interaction with RasG12D-GTP is further from an evolutionary optimum than its interaction with RasWT-GTP (Fig. 2A top left and bottom left). While there is a moderate positive correlation between single-mutation enrichments in the HI gate (R = 0.53), several Nore1A mutations display an anticorrelated pattern, showing positive enrichment for RasG12D-GTP but negative enrichment for RasWT-GTP (Fig. 2G). This suggests the presence of variants that preferentially bind RasG12D-GTP over RasWT-GTP (Fig. 2G, upper left quadrant). Among those, the most frequent mutations occur at Nore1A positions 218 and 221 that interact with switch II on Ras and 277 that interacts with switch I (Figure 1A). In contrast, very few single mutations preferentially favor RasWT-GTP over RasG12D-GTP (Fig. 2G, lower right quadrant). RasG12D-GDP also shows a higher number of affinity-enhancing single mutations compared to that of RasWT-GDP, specifically at positions 218 and 277 (Fig. 2A, bottom right). Analysis of double mutants reveals a large number of variants with specificity toward RasG12D-GTP (Fig. 2H, upper left quadrant), as well as a smaller subset that preferentially bind RasWT-GTP (Fig. 2H, lower right quadrant). In summary, both single and double mutations in Nore1A can sense the oncogenic mutation on Ras, with a number of mutations in Nore1A likely increasing affinity for RasG12D but not for RasWT.

### Differences in the Nore1A-RasWT and RasG12D binding landscapes could be explained by differences in complex dynamics

Since the oncogenic Ras mutation site is not located at the direct Nore1A–Ras binding interface (Fig. 1A), the differences in the binding landscapes are unlikely to result from direct interactions involving residue 12. Instead, they may reflect changes in the dynamics of the complexes. To test this hypothesis, we performed molecular dynamics (MD) simulations of complexes containing either RasWT-GTP or RasG12D-GTP paired with Nore1A WT or selected affinity-enhancing Nore1A variant (Movies S1-S4 in SI Appendix). Using the MD trajectory files, we compared the resulting complex dynamics by calculating the Root Mean Square Fluctuation (RMSF) for each residue of both Ras and Nore1A (Fig. 3). Our data show that, in complexes with Nore1A WT, RasG12D exhibits overall higher RMSF values than RasWT, indicating increased flexibility across several regions, including the vicinity of the mutation site, switch I, and the α3-loop (Fig. 3A), consistent with previous MD simulations of unbound RasWT and RasG12D (35), suggesting that this intrinsic dynamic property of RasG12D is retained upon effector binding. In contrast, Nore1A displayed largely similar RMSF profiles when bound to RasWT or RasG12D, with only a modest increase in flexibility in the α1 region when bound to RasG12D.

**Fig. 3.**
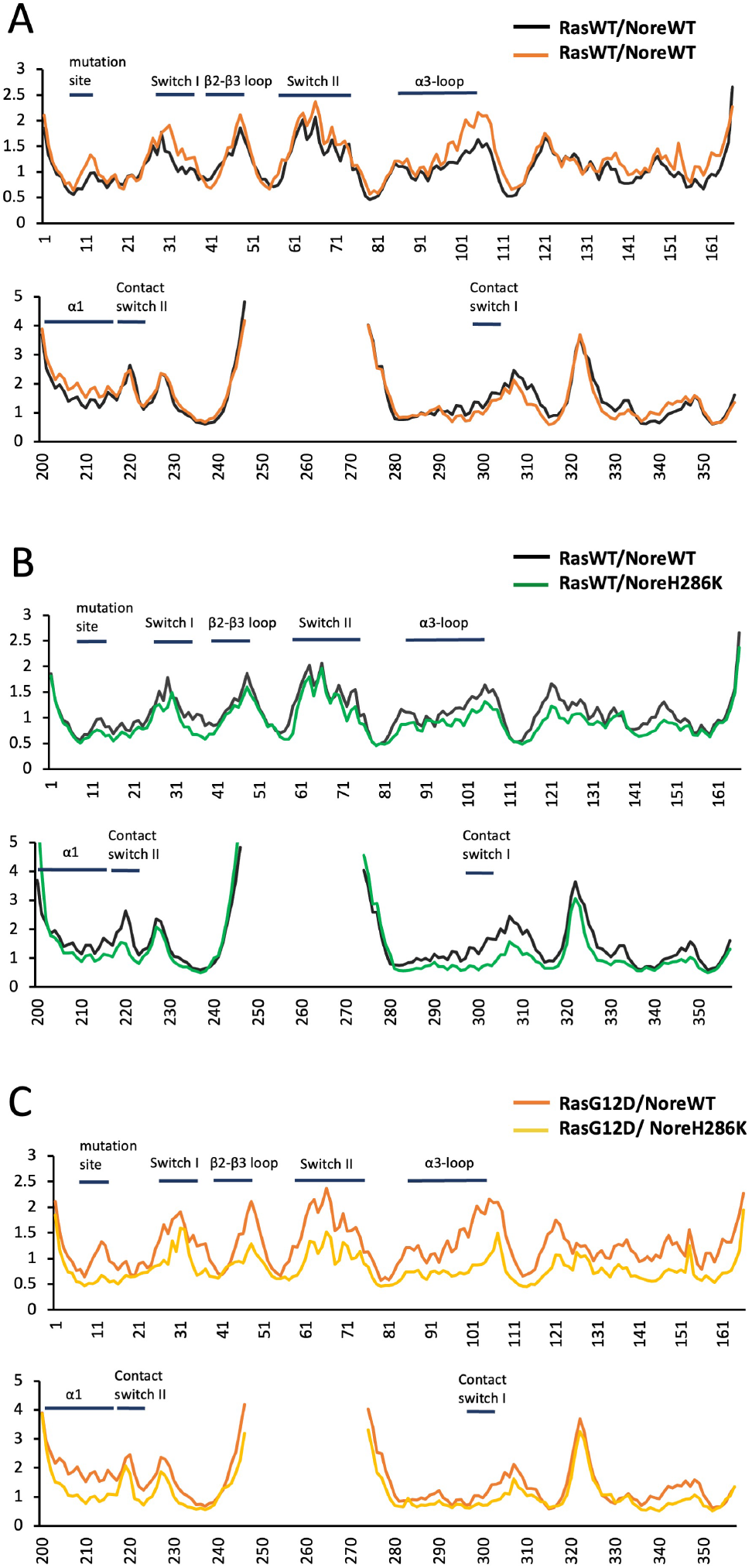
(A) RMSF of Ras/Nore1A variant complexes. Pairwise comparison of two mutant complexes: (A) Nore1A WT interacting with RasWT and RasG12D; (B) Nore1A WT and Nore1A H286K interacting with RasWT; (C) Nore1A WT and Nore1A H286K interacting with RasG12D. During the simulations, the long, unstructured loop in Nore1A (residues 249–273) was replaced with a shorter, eight-residue loop connecting P248 to S274 to prevent excessive loop movements, as described in the Methods. RMSF values were calculated from MD simulations of the complexes performed under identical conditions. Key regions of Ras and Nore1A are indicated above the RMSF curves.

Introducing the affinity-enhancing H286K mutation in Nore1A resulted in a moderate reduction in flexibility throughout RasWT and in the Nore1A regions contacting RasWT switch I and II ((Fig. 3B). When RasG12D was simulated with the same Nore1A mutant, RMSF values were markedly reduced across RasG12D, indicating a substantial decrease in its flexibility (Fig. 3C. Reduced flexibility was also observed in specific regions of Nore1A, including the α1 region and the regions contacting switch I and II on Ras. Similarly, large decrease in RMSF over RasG12D residues was observed when in complex with another affinity-enhancing mutation on Nore1A, Q284T (Figure S4 in SI Appendix). Together, these results suggest that the Nore1A affinity-enhancing mutations strengthen binding to Ras variants by reducing the flexibility of both proteins. This effect is more pronounced in complexes with RasG12D than with RasWT, consistent with the intrinsically higher flexibility of RasG12D.

### Building quantitative binding landscapes for Nore1A-Ras interactions

To move beyond the qualitative information provided by the enrichment heat maps, we used the enrichment values of each mutant across four affinity gates to predict ΔΔ*G*_bind_. Specifically, we applied a methodology previously developed by our group that uses a small dataset of experimental ΔΔ*G*_bind_ values, measured by biophysical techniques on purified proteins, to normalize enrichment values and derive ΔΔ*G*_bind_ values for all mutations detected in the NGS experiment (3, 33). To establish an experimental dataset for normalization, we expressed and purified 16 Nore1A variants spanning the full range of enrichment values in the HI, WT, SL, and LO gates for RasWT-GTP, from strongly positive to strongly negative. We measured the binding affinities of these variants for Ras using a Guanine nucleotide Dissociation Inhibitor (GDI) assay, a widely used method for determining binding affinity (*K*_*D*_) values of Ras–effector interactions (9, 12). The GDI assay is based on the principle that effector binding to Ras slows down the dissociation of a fluorescent GTP analogue, Mant-GTP (2*′*/3*′*-O-(N-methyl-anthraniloyl)-guanosine 5*′*-triphosphate) from Ras, enabling determination of the Ras interaction’s binding affinity for a particular Nore1A mutant (Fig. 4A).

**Fig. 4.**
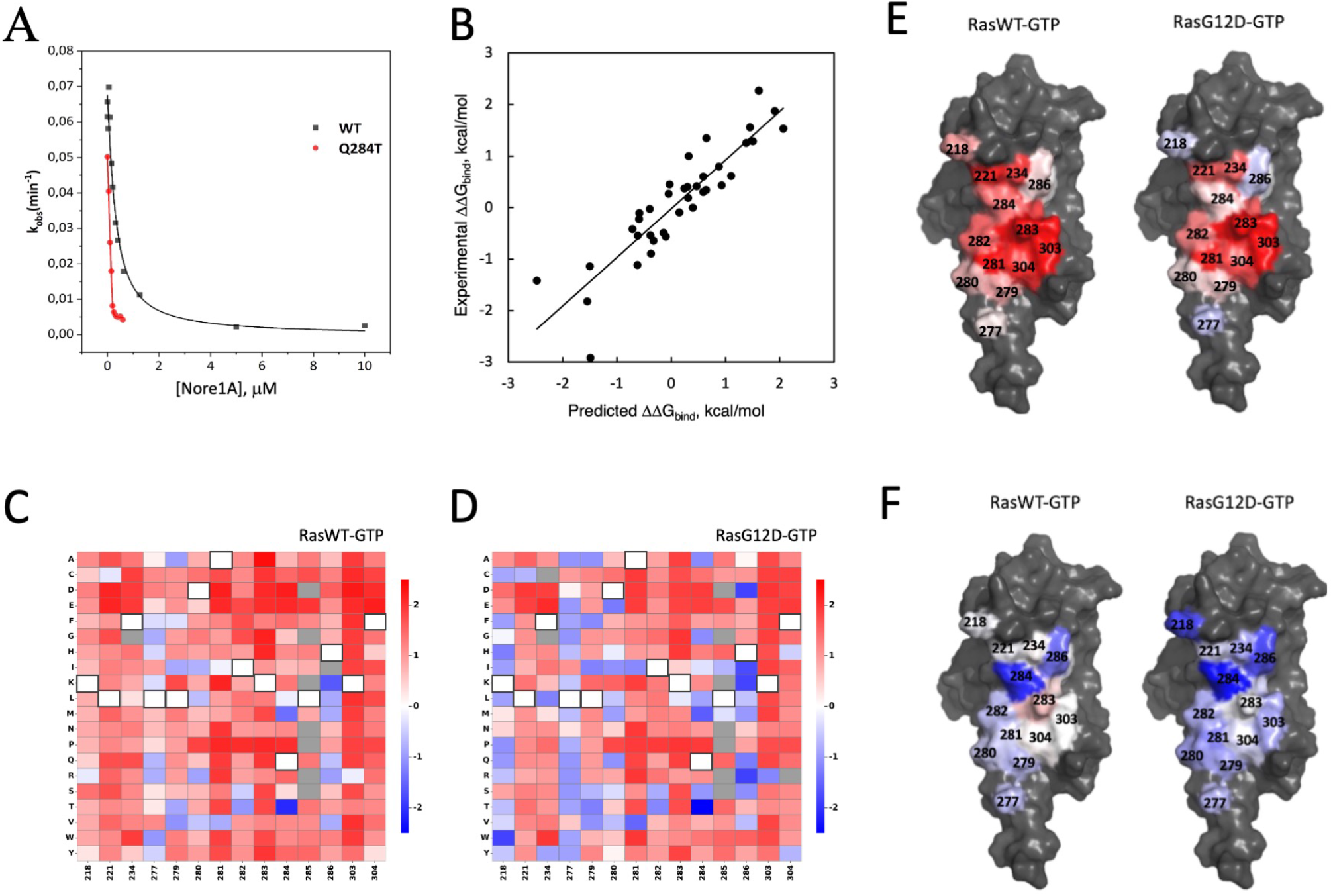
(A) GDI assays showing binding of Nore1A WT and Nore1A Q284T to RasWT-GTP, used to determine *K*_*D*_ values for selected Nore1A mutants. (B) Correlation between experimentally measured and predicted ΔΔ*G*_bind_ values for purified Nore1A mutants binding to RasWT-GTP and RasG12D-GTP. Predictions were obtained using leave-one-out cross-validation. The solid line represents a linear fit to the data, with a Q-value of 0.90 and a standard deviation (*σ*) of 0.42 kcal/mol. (C–D) Quantitative binding landscapes for all single mutations in the Nore1A/RasWT-GTP complex (C) and Nore1A/RasG12D-GTP complex (D). The y-axis shows amino acid identity, and the x-axis shows position in Nore1A. Mutations are color-coded from highly stabilizing (blue) to highly destabilizing (red). Wild-type amino acids are shown in white squares, and mutations for which no predictions are available are shown in gray. (E) Average per-position ΔΔ*G*_bind_ values mapped onto the structures of Nore1A interacting with RasWT-GTP (left) and RasG12D-GTP (right). Values range from −1.0 kcal/mol (blue) to +1.0 kcal/mol (red). (F) Minimum per-position ΔΔ*G*_bind_ values mapped onto the structures of Nore1A interacting with RasWT-GTP (left) and RasG12D-GTP (right). Values range from −2.0 kcal/mol (blue) to +2.0 kcal/mol (red).

Our experiments showed that WT Nore1A binds to RasWT-GTP with slightly higher affinity than to RasG12D-GTP, with measured *K*_*D*_ values of 0.22 and 0.55 µM, respectively. We then measured the *K*_*D*_ values of Nore1A variants in complex with RasWT-GTP and RasG12D-GTP and calculated the corresponding changes in binding free energy (ΔΔ*G*_bind_) relative to that of WT Nore1A (Table S1 in SI Appendix). We further supplemented our dataset with ΔΔ*G*_bind_ measurements for Nore1A variants interacting with RasWT-GTP, reported in a previous study that employed the same experimental method (24), yielding a combined experimental dataset of 38 ΔΔ*G*_bind_ measurements. We also attempted to measure the binding affinities of these Nore1A variants for Ras in the GDP-bound state. However, these measurements required high concentrations of Nore1A variants, which tended to aggregate at concentrations above 10 µM. We therefore restricted the experimental dataset to interactions with GTP-bound Ras.

This experimental dataset was used to build a model that converts the four enrichment values for each Nore1A mutant into a ΔΔ*G*_bind_ value. We explored a range of model architectures, including simple linear regressions on the enrichment scores, models augmented with nonlinear terms, and ensemble tree-based methods. We evaluated different models by their ability to exhibit high correlation with this experimental dataset in a leave-one-out cross-validation test and to be able to extrapolate to further areas outside the range. Our best-performing model was based on a modified Extremely Randomized Trees algorithm that incorporated the identity of the affinity gate with the maximum enrichment. The model predictions showed strong agreement with ΔΔ*G*_bind_ values measured experimentally on purified proteins (Fig. 4B; *Q*^2^ = 0.81; RMSE = 0.42 kcal/mol in leave-one-out cross-validation test). The trained model was subsequently applied to predict ΔΔ*G*_bind_ values for all Nore1A single and double mutants detected by NGS, generating quantitative binding landscapes for Nore1A interactions with Ras i n four distinct states: RasWT-GTP, RasG12D-GTP, RasWT-GDP, and RasG12D-GDP. To avoid possible artifacts, in our predictions we excluded mutations detected in fewer than three affinity gates or exhibiting inconsistent enrichment patterns across the gates (see Methods). This filtering yielded coverage of 97% of all possible single mutations in our library and 40–50% of all possible double mutations. The resulting binding landscapes are presented in Fig. 4C and D for single mutations in the RasWT-GTP and RasG12D-GTP states, respectively, and in Figures S5 and S6 in SI Appendix for single mutations in the Ras-GDP states and for all double mutations, respectively. The binding landscapes were then analyzed to characterize the diverse effects of mutations on binding affinity and interaction specificity.

### Structural signature of the binding landscapes

First, we analyzed how the ΔΔ*G*_bind_ values are distributed across different positions on Nore1A, revealing the structural signature of the Nore1A-Ras binding landscapes (Fig. 4E and F). Fig. 4E shows the mean ΔΔ*G*_bind_ value across 19 mutations calculated for each Nore1A binding interface position. This figure reveals clear shared hotspots for both RasG12D–GTP and RasWT–GTP complexes, centered in the core of the interface, particularly at positions 283 and 303 that both form important salt bridges to Ras and nearby positions 281 and 304 occupied by hydrophobic amino acids. The Nore1A interface with RasWT-GTP lacks prominent cold-spot positions at the periphery of the interface, with nearly all positions exhibiting, on average, a destabilizing effect on binding, whereas its interface with RasG12D-GTP contains several cold-spot positions at the periphery (Fig. 4E, shown in blue). A similar difference between the Nore1A-RasG12D and Nore1A-RasWT interfaces is observed when comparing the minimum ΔΔ*G*_bind_ values across Nore1A positions (Fig. 4F). The strongest affinity-enhancing mutations are concentrated in the interface core, at positions 284 and 286, in both complexes. However, in the Nore1A/RasWT–GTP interface, affinity-enhancing mutations are absent at nearly half of the interface positions. In contrast, in the Nore1A/RasG12D–GTP interface, affinity-enhancing mutations are observed at all sites except for position 283. The analysis of Nore1A interactions with Ras in the GDP state revealed a highly similar structural distribution of hotspots and cold spots across the interface (Figures S7E in SI appendix). Interestingly, the hotspot region, comprising positions 281, 283, 303, and 304, is conserved across all four Ras isoforms studied here. Remarkably, this hotspot region directly contacts a hotspot region on Ras, particularly residues D38 and Y40, which have been identified as key energetic residues in Ras interactions with Raf and other effectors (32). Collectively, these findings indicate that the overall energetic architecture of the Nore1A/RasWT and Nore1A/RasG12D binding interfaces is structurally conserved and largely independent of nucleotide state. However, the Nore1A/RasG12D–GTP interface contains a greater number of non-optimal positions, resulting in a greater potential for affinity enhancement in this complex.

### Binding Energetics of Single and Double Mutations

Next, we analyzed the distribution of ΔΔ*G*_bind_ values for Nore1A mutants interacting with RasWT–GTP and RasG12D–GTP (Fig. 5A-D). Single mutations in Nore1A exhibited ΔΔ*G*_bind_ values ranging from −1.8 to +2.3 kcal/mol when interacting with RasWT–GTP, with most mutations being moderately destabilizing and an average ΔΔ*G*_bind_ of 0.69 kcal/mol. For Nore1A interactions with RasG12D–GTP, ΔΔ*G*_bind_ values ranged from −2.9 to +1.8 kcal/mol, with an average of 0.44 kcal/mol. Thus, the distribution for RasG12D–GTP is shifted toward more negative ΔΔ*G*_bind_ values relative to those of RasWT–GTP, indicating that the Nore1A interaction with the oncogenic Ras variant is less optimized. Next, we analyzed the distribution of ΔΔ*G*_bind_ values for double mutations. Interestingly, these distributions closely resembled those observed for single mutations, spanning −2.9 to +2.2 kcal/mol for RasWT–GTP and RasG12D–GTP. Notably, we did not observe more extreme ΔΔ*G*_bind_ values that might be expected from additive effects of two affinity-enhancing or two strongly destabilizing mutations. The average ΔΔ*G*_bind_ values were 0.90 and 0.80 kcal/mol for RasWT–GTP and RasG12D–GTP, respectively, which are slightly higher than the corresponding averages for single mutations. To further understand how the effects of two single mutations combine to produce the effect of a double mutation, we examined in detail all double mutations involving positions 234 and 284, which are close in space but distant in sequence (Fig. 5E and F). Our analysis shows that most double mutants containing the affinity-enhancing Q284T mutation are also affinity-enhancing, regardless of the identity of the second mutation at position 234, in both complexes (blue vertical lines in Fig. 4E and F). Similarly, double mutants containing highly destabilizing mutations at position 284 (D, E, and P) are generally destabilizing regardless of the mutation at position 234 (red vertical lines). Highly destabilizing mutations at position 234 (D, E, and W) similarly are destabilizing across most of mutations at position 284 and appear as horizontal red lines. These results support the validity of the beneficial and deleterious single mutations identified by the NGS data, suggesting that they introduce or disrupt specific molecular interactions that are preserved in most double mutants and are frequently observed in both complexes. However, beneficial or disruptive interactions introduced by two new individual mutations cannot always be accommodated simultaneously, resulting in mutational effects that are smaller than additive. To further analyze the additivity of mutation effects throughout our extensive quantitative datasets of ΔΔ*G*_bind_ values, we calculated the interaction coupling energy between the two mutations Δ*G*_*i*_ (see Methods) in complexes where Nore1A interacts with RasWT-GTP and RasG12D-GTP. We classified the observed mutation pairs into three categories: additive, exhibiting positive and negative epistasis, based on the magnitude of the coupling energy, Δ*G*_*i*_ and plotted the additive/epistatic properties of each mutation at each position (Figure S8 in SI Appendix). Our results showed that additivity of two mutations depends not only on the pairs of positions but also on the identity of the mutation. Furthermore, the coupling energy analysis revealed that, for the Nore1A–RasWT–GTP interaction, 58% of mutations were additive, 40% exhibited positive epistasis, and only 2.3% displayed negative epistasis. In contrast, the Nore1A–RasG12D–GTP interaction showed a comparable percentage of additive mutations of 56%, but an altered distribution of epistatic effects, with 29% of mutations exhibiting positive epistasis and 15% exhibiting negative epistasis. A similar relationship between interface optimization level and epistasis was observed in our previous studies of three homologous serine protease–BPTI complexes where the most optimal bovine trypsin–BPTI complex exhibited the greatest proportion of positive epistasis, whereas the lowest-affinity mesotrypsin–BPTI complex showed the highest proportion of negative epistasis (3). Thus, mutational coupling results highlight the evolutionary differences between the Nore1A–RasWT–GTP and the Nore1A–RasG12D–GTP complexes and support greater optimization of the Nore1A interface with WT Ras in comparison to its oncogenic mutant.

**Fig. 5.**
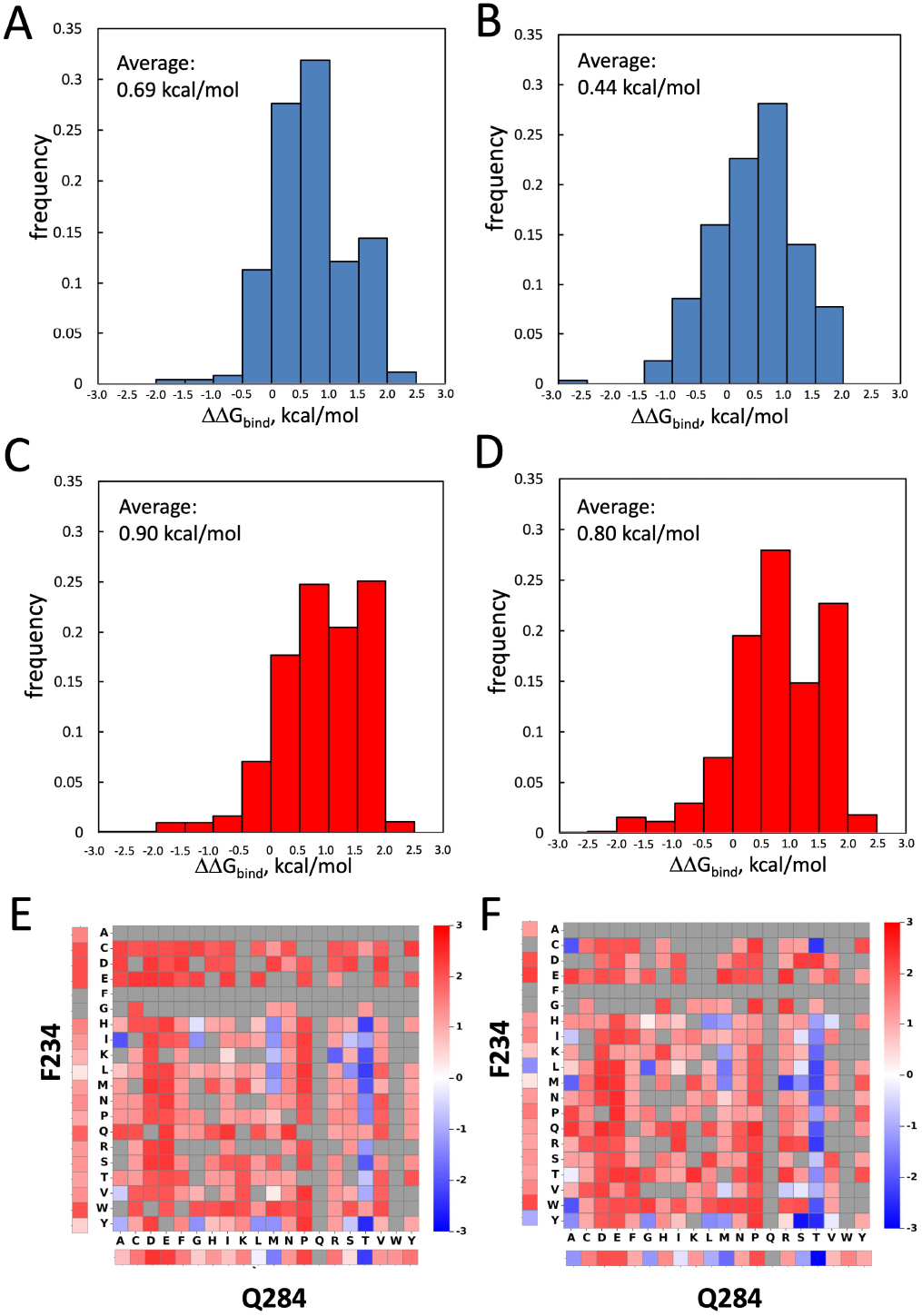
(A)-(D) Histograms of the predicted ΔΔ*G*_bind_ values for all detected single and double mutations. (A) Single mutations for the Nore1A-RasWT-GTP interaction; (B) Single mutations for the Nore1A-RasG12D-GTP interactions; (C) Double mutations for the Nore1A-RasWT-GTP interaction; (D) Double mutations for the Nore1A-RasG12D-GTP interaction. ΔΔ*G*_bind_ values for all double mutations that involve positions 234 and 284 for the Nore1A-RasWT-GTP interaction (E) and for the Nore1A-RasG12D-GTP interaction (F) colored from red (destabilizing) to blue (stabilizing). Grey squares represent mutations that have not been detected. On the left and at the bottom of each graph the ΔΔ*G*_bind_ values for the single mutations at position 234 and 284 are given, respectively.

## Discussion

In this study, we explored the binding energy landscape of the Ras effector Nore1A in complex with Ras, a molecular switch that cycles between active GTP-bound and inactive GDP-bound states. To our knowledge, this is the first comprehensive study to examine how mutations in a Ras effector alter its binding affinity for both nucleotide states of Ras, thereby defining the potential signalling output of the interaction. Systematically mapping these effects revealed that the Nore1A/RasWT-GTP binding landscape is evolutionarily near-optimal: only a small number of single mutations increased its affinity, and most changes were modest, with effects not exceeding 2 kcal/mol. While a high degree of evolutionary optimization characterizes many ultra-high-affinity enzyme–inhibitor complexes (4, 33), such optimization was unexpected for a medium-affinity signaling complex such as Nore1A–Ras. One possible explanation for the high optimality of the Nore1A/RasWT-GTP interface is its flat shape and the abundance of polar intermolecular interactions that are, for the most part, already satisfied. Most polar residues on Ras are already engaged in intermolecular salt bridges and hydrogen bonds with Nore1A, leaving relatively few opportunities to introduce new favorable interactions.

The high degree of optimization observed for Nore1A–RasWT-GTP is partially lost upon introduction of the oncogenic G12D mutation into Ras. Although the overall structural and energetic signatures of the Nore1A–RasG12D interface remain largely unchanged, the G12D complex contains a substantially larger number of affinity-enhancing mutations. Our MD simulations suggest that this loss of optimality may arise from the greater flexibility of RasG12D relative to RasWT, creating opportunities for Nore1A mutations that reduce Ras flexibility and thereby enhance binding affinity. This interpretation is consistent with previous MD simulations of RasG12D alone showing that the G12D mutation increases flexibility in the Switch II region of Ras (35), as well as with NMR studies demonstrating a greater population of Switch I in the inactive conformation in RasG12D compared to RasWT (36). Thus, Nore1A mutations that stabilize these dynamic regions may enhance binding by constraining RasG12D toward an effector-competent conformation. Consistent with this model, the H286K mutation in Nore1A decreases Ras flexibility and substantially increases affinity for RasG12D. The high degree of optimization observed for Nore1A–RasWT-GTP is only slightly reduced in the Nore1A–RasWT-GDP complex, despite the substantial decrease in affinity following GTP hydrolysis. Although the low-affinity Nore1A/Ras–GDP interaction might be expected to contain numerous cold spots—positions amenable to affinity-enhancing mutations—we observed no such enrichment. Instead, the energetic and structural signatures across the interface remained largely unchanged relative to those of the higher-affinity Nore1A/Ras–GTP complex. Thus, the lower affinity of Nore1A for Ras-GDP does not appear to reflect widespread under-optimization of the interface, but rather the nucleotide-dependent conformational state of Ras.

Previous studies showed that single and double mutations in the Ras effector Raf can selectively increase its affinity for Ras-GDP, thereby narrowing the affinity gap between the GDP-and GTP-bound states (9, 37). Structural analysis of one such Raf double mutant in complex with Ras-GDP revealed that Switch I of Ras in this complex adopts a GTP-like conformation, the only conformation compatible with effector binding. However, in this structure, Switch I retains residual flexibility, limiting the affinity of the Raf mutant for Ras-GDP and thus preserving Ras switching properties. In our study, we similarly identified a small number of single and double mutations in Nore1A that narrow the affinity gap between Ras-GTP and Ras-GDP. Most of these mutations occur at positions that interact directly with the two Ras switch regions, including K218I/V/L and H286K/R, which interact with Switch II, I282T, which interacts with Switch I and L277Y and L279V that form intermolecular beta-sheet with Ras. Notably, we did not identify any single or double mutations that completely reversed the binding preference of Nore1A from Ras-GTP to Ras-GDP, as this would require the same Nore1A mutant to exhibit ΔΔ*G*_bind_ more negative by at least 4 kcal/mol for Ras-GDP compared to Ras-GTP. Together, these findings suggest that, within the architecture of the Nore1A–Ras interface, mutations can reduce but not eliminate the intrinsic preference for Ras-GTP. This constraint may represent an evolutionary safeguard that preserves the function of Ras as a molecular switch by preventing mutations in effector proteins from readily converting them into preferential binders of the inactive GDP-bound state.

Nore1A is a unique effector that counterbalances the proliferative signaling mediated by other Ras effectors under normal physiological conditions. In cancer, when Ras signaling becomes dysregulated, Nore1A does not acquire recurrent mutations but is instead frequently inactivated through epigenetic mechanisms, particularly promoter hypermethylation. Such epigenetic silencing may be particularly advantageous for tumor cells because disrupting Nore1A function through mutations in its Rasbinding interface would be relatively inefficient, given that Nore1A has one of the highest affinities and lowest dissociation rates among Ras effectors. Indeed, mutations across the Nore1A interface produce, on average, only modest destabilization of the complex ( 0.8 kcal/mol), with a maximum effect of 1.8 kcal/mol, which may be insufficient to eliminate the signaling that drives Nore1A-mediated senescence and apoptosis. This may be particularly important in the context of oncogenic Ras, where the RasG12D binding landscape contains a relatively large number of mutations capable of further enhancing Nore1A affinity. Thus, rather than disrupting the Nore1A–Ras interaction through interface mutations, epigenetic silencing may provide a more effective means of eliminating Nore1A signaling and preventing this tumor-suppressive pathway from counterbalancing oncogenic Ras signaling.

Beyond providing insight into the evolutionary and energetic determinants of Ras–effector recognition, our findings have implications for the design of selective Ras protein-based inhibitors. The distinct binding landscapes of Nore1A-RasWT and Nore1A-RasG12D demonstrate that an oncogenic mutation can create opportunities for engineering interaction partners with altered specificity, even when the mutation does not directly contact the binding interface. We identified several such mutations, particularly at Nore1A positions 218, 277, and 286 that could selectively enhance Nore1A affinity for RasG12D over RasWT. We thus proved that systematic mapping of binding landscapes can facilitate the design of selective Nore1A variants for oncogenic Ras mutants. These protein engineering efforts could complement small-molecule approaches that target the nucleotide-binding pocket or allosteric sites on Ras by exploiting changes in protein dynamics and energetic coupling that are not necessarily accessible through conventional drug-binding pockets. More broadly, our findings demonstrate how mutational binding landscapes can reveal exploitable differences between oncogenic and wild-type proteins and provide a framework for designing selective protein-based inhibitors against disease-associated Ras variants.

## Materials and Methods

### Construction of the Nore1A library

Fourteen positions on Nore1A at the direct binding interface with Ras were selected for site-saturation mutagenesis (19 substitutions per position), generating both single- and double-mutant libraries (Twist Bioscience). Amplified library DNA was cloned into the pCTCON2 yeast surface display vector by homologous recombination in *S. cerevisiae* strain EBY100, enabling Aga2p-fusion surface display with a C-terminal myc tag. Full cloning and transformation details are provided in SI Methods.

### Protein expression, purification, and nucleotide loading

The HRas and HRasG12D G-domains (residues 1–261) were expressed in *E. coli* BL21(DE3) as His-tagged, AviTag-biotinylatable constructs and purified by Ni2+-affinity and size-exclusion chromatography, followed by TEV cleavage of the His-tag (final yield 6.7 mg purified

Ras per liter of culture). Purified Ras was then loaded with GDP or the nonhydrolyzable GTP analog GppNHp by EDTA-mediated Mg2+ chelation and molar-excess nucleotide incubation, as previously described (38); excess nucleotide was removed by desalting, and loaded protein was flash-frozen for storage. Full expression, purification, and nucleotide-exchange conditions are provided in SI Methods.

### FACS sorting and NGS analysis

Yeast libraries displaying Nore1A variants were incubated with Ras-GTP, Ras-GDP, RasG12D-GTP, or RasG12D-GDP at concentrations optimized per condition, and sorted by FACS (BD FACSAria III) into four affinity-based populations (HI, WT, SL, LO) relative to a WT-defined reference gate, following Chao et al. (39). Plasmid DNA recovered from each sorted population was sequenced in duplicate (Illumina MiSeq/NextSeq, ≥ 106 reads/sample). Reads were merged and quality-filtered (FLASH, FastQC), and single and double mutants were identified and counted using a custom MATLAB pipeline, discarding reads with more than two mutations and variants occurring fewer than 10 times in the naïve library. Full sorting parameters, media, and library preparation for NGS are provided in SI Methods.

### Enrichment calculation and ΔΔ*G*_bind_ prediction

For each NoreA variant sequence, the number of occurrences was counted in both the sorted and the naïve populations, and its frequency (*f*_*i*_) in the particular gate was calculated (Eq. 1).

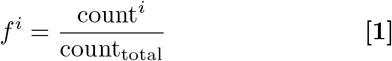

Using the relative frequencies in a sorted population and the naïve library, the enrichment (*e*_*i*_) of each NoreA mutant was then computed (Eq. 2):

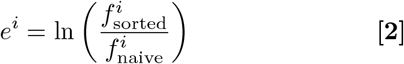

The enrichment values were further normalized in each gate to range from -1 to +1 using the the maximum and minimum enrichment values, *e*^max^ and *e*^min^, respectively (Eq. 3):

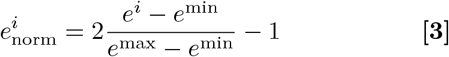

Variants with enrichment values in fewer than 3 of 4 gates were excluded. ΔΔ*G*_bind_ was predicted from the fourgate enrichment profile using a state-aware ensemble of Extremely Randomized Trees regressors (40, 41), trained on 38 experimental ΔΔ*G*_bind_ values (Table S1) and validated by leave-one-out cross-validation. The trained model was applied to all detected single and double mutants for all four Ras/nucleotide states. The coupling energy Δ*G*_int_ upon two mutations X and Y was calculated according to:

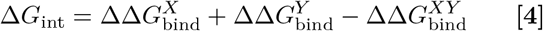

with significance thresholds defined by propagated prediction uncertainty. Full model architecture and feature details in SI Methods.

### Nore1A mutant construction and *K*_*d*_ measurement

Selected Nore1A mutants (Table S1 in SI appendix) were generated by Transfer-PCR (42), expressed and purified from BL21(DE3), and their binding affinity (*K*_*d*_) for RasWT-GTP and RasG12D-GTP was measured by a guanine nucleotide dissociation inhibition (GDI) assay using mant-labeled nucleotides and fit to a quadratic binding model to calculate *K*_*d*_ (43). Full assay conditions and fitting equations in SI Methods.

### Molecular dynamics simulations

MD simulations of the Ras–Nore1A complex (PDB 3DDC) were performed in OpenMM (44) using the AMBER14 force field with TIP3P-FB water, PME electrostatics, and a Langevin integrator at 300 K. A 50-ns production run followed energy minimization and equilibration; trajectory stability was confirmed by RMSD, and per-residue flexibility was assessed by RMSF. Full simulation parameters in SI Methods.

## Data, Materials, and Software Availability

All the data are included in the Supporting Information.

## ACKNOWLEDGMENTS

We thank Gideon Schreiber (WIS) for comments on our paper. This work was supported by the Israel Science Foundation grant 3486/20, ICRF, and the University of Toronto/HUJI Research alliance. We also acknowledge support from the NSF/BSF grant 2022685 and NIH R01CA 258274.

