## Supplementary material for "The binding landscape of Ras/effector interactions provides insights on Ras switching and oncogenesis": SI Appendix

### Extended Methods & Materials

#### Construction of the Nore1A library

Fourteen positions on Nore1A that are in direct binding interface with Ras were selected for randomization. This library was synthesized using solid-phase DNA synthesis technology provided by Twist Bioscience (San Francisco, CA, USA). Each of the selected positions was individually subjected to site-saturation mutagenesis, replacing the native codon with codons encoding 19 canonical amino acids, thereby generating a comprehensive single-mutant library. To complement the single-substitution library, a combinatorial library of double mutants was also constructed. Each site-variant sub-library was received as 50 ng of lyophilized DNA per vial. To obtain sufficient quantities for downstream applications, the DNA was amplified by polymerase chain reaction (PCR). Reactions were performed using primer pairs designed to amplify the full-length gene with flanking regions for homologous recombination. These primers were ordered from Integrated DNA Technologies (IDT) with the following sequences: Nore_Forward: 5′-GCT CTG GTG GAG GCG GTA GCG GAG GCG GAG-3′, Nore_Reverse: 5′-CTA TTA CAA GTC CTC TTC AGA AAT AAG CTT TTG TTC GGA TCC-3′. The PCR reaction contained 1 μL of Phusion DNA Polymerase (New England Biolabs), 20 μL of 5X buffer, 2 μL of 10 mM dNTPs, 2.5 μL of 10 μmol each primer and 60 μL of water, 12 μL of template. PCR conditions were as follows: initial denaturation at 98 °C for 30 s, 12 denaturation cycles at 98 °C for 10 s, annealing at 66 °C for 30 s, extension at 72 °C for 30 s and a final extension at 72 °C for 10 min. Following amplification, PCR products were purified using the QIAquick PCR Purification Kit (QIAGEN), according to the manufacturer's instructions. DNA concentrations were measured using a NanoDrop spectrophotometer (Thermo Scientific). The typical yield after amplification was approximately 10 μg per sublibrary. The purified PCR fragments were cloned into the pCTCON2 yeast surface display vector, which enables N-terminal fusion of the target protein to Aga2p for presentation on the surface of *Saccharomyces cerevisiae*, and appends a C-terminal myc epitope tag for detection and quantification of surface expression by flow cytometry. Cloning was carried out *in vivo* by homologous recombination in *S. cerevisiae* through co-transformation of the PCR-amplified inserts and linearized vector DNA. The pCTCON2 vector was linearized with NheI and BamHI, generating appropriate flanking regions of homology to promote efficient recombination and insertion of the target sequence. We prepared four parallel transformations. A total of 5 μg of linearized pCTCON2 vector and 20 μg of library DNA (amplified separately in 15 vials and subsequently combined) were co-transformed into *S. cerevisiae* strain EBY100 using the standard lithium acetate/PEG transformation protocol. Electroporation was performed using a MicroPulser™ Electroporation Apparatus (Bio-Rad), following the manufacturer's guidelines and established protocols for yeast surface display. DNA samples with high salt concentrations were treated with Pellet Paint® Co-Precipitant (Merck) to remove excess salt prior to transformation.

**Protein expression and purification**

The HRas and HRasG12D genes corresponding to the G domain (residues 1-261) were cloned into a modified pET28a expression vector containing an N-terminal TEV protease cleavage site (ENLYFQ/G) and a C-terminal AviTag™ sequence (GLNDIFEAQKIEWHE) to allow in vivo biotinylation. The resulting construct was transformed into *Escherichia coli* BL21(DE3) cells co-transformed with the BirA expression plasmid (pACYC184 vector encoding the birA gene) for biotin ligase production. Cells were cultured in Luria-Bertani (LB) medium supplemented with kanamycin (50 µg/mL) and chloramphenicol (34 µg/mL) at 37 °C with shaking at 225 rpm. Upon reaching an optical density at 600 nm ($OD_{600}$) of ~0.8, protein expression was induced with 0.1 mM IPTG , and incubation was continued for an additional 3 hours. Cells were harvested by centrifugation at 5,000 × g for 30 minutes at 4 °C, then resuspended in lysis buffer (20 mM Tris-HCl, 500 mM NaCl, 5 mM $MgCl_2$, pH 7.5) supplemented with a protease inhibitor cocktail (cOmplete™ Mini, EDTA-free, Roche). Cell lysis was performed by sonication (9 seconds ON / 9 seconds OFF, 40% amplitude) for a total of 5 minutes. The lysate was clarified by centrifugation at 20,000 × g for 40 minutes at 4 °C. The supernatant was incubated with $Ni^{2+}$-NTA agarose resin (PureCube, Cube Biotech) pre-equilibrated with Buffer A(20 mM Tris-HCl, 100 mM NaCl, 5 mM $MgCl_2$, 5 mM imidazole, pH 7.5) for 1 hour at 4 °C. The resin was loaded onto a gravity-flow column and washed sequentially with at least 20 column volumes (CV) of Buffer A containing increasing concentrations of imidazole (20 mM, 50 mM, 200 mM, and 500 mM). The Ras protein was eluted with 200 mM imidazole, and 1 mM DTT was added to each fraction to prevent precipitation during storage. Eluted protein fractions were pooled, concentrated, and further purified by size-exclusion chromatography (SEC) using a Superdex 75 Increase 10/300 GL column (Cytiva) equilibrated with Buffer A. The peak fractions corresponding to monomeric Ras were collected. To remove the N-terminal $His_6$-tag, the pooled protein was treated with TEV protease at a ratio of 30–60 µg per 1 mg of Ras protein and incubated overnight at 4 °C. The digestion mixture was passed over $Ni^{2+}$-NTA resin to remove the cleaved His-tag, uncleaved protein, and TEV protease. The flow-through, containing the tag-free Ras protein, was collected and concentrated using Amicon Ultra centrifugal filters with a 3 kDa molecular weight cutoff (Millipore). The cleaved Ras protein was further purified by size-exclusion chromatography using a Superdex 75 column (GE Healthcare) equilibrated in buffer suitable for downstream applications (20 mM Tris-HCl pH 8.0, 150 mM NaCl, 1 mM DTT). The final yield of purified Ras protein was approximately 6.7 mg per liter of bacterial culture.

**Nucleotide Exchange**

Recombinant Ras protein was subjected to buffer exchange prior to nucleotide loading to reduce $MgCl_2$ concentration. The storage buffer (50 mM Tris, pH 7.5, 100 mM NaCl, 5 mM $MgCl_2$) was exchanged to 50 mM Tris (pH 7.5), 100 mM NaCl, and 0.5 mM $MgCl_2$ using Amicon Ultra centrifugal concentrators (3-kDa molecular weight cutoff). All buffer exchange steps were performed at 4 °C. The procedure was adapted from a previously described guanine nucleotide exchange assay (1). Ras nucleotide exchange was performed by incubating protein with a 20-fold molar excess of GDP or GppNHp (a nonhydrolyzable GTP analog) in exchange buffer containing 20 mM Tris (pH 7.5), 100 mM NaCl, 0.5 mM $MgCl_2$, 10 mM EDTA, and 3 mM DTT for 1 h at 4 °C. EDTA chelates $Mg^{2+}$ ions that coordinate with the β and γ phosphates of GTP (or the β phosphate of GDP), thereby destabilizing nucleotide binding and facilitating exchange. A control reaction without added nucleotide was performed in parallel. Reactions were terminated by addition of $MgCl_2$ to a final concentration of 10 mM, followed by incubation for 30 min at 4 °C to stabilize nucleotide binding. Excess nucleotide was removed using Zeba Spin Desalting Columns (Thermo Fisher Scientific) according to the manufacturer's instructions. Protein concentration was determined using a bicinchoninic acid (BCA) assay with bovine serum albumin (BSA) as standard. The final yield was approximately 4 mg of nucleotide-loaded Ras per liter of bacterial culture. Samples were supplemented with 7.5% glycerol, aliquoted, flash-frozen, and stored at −80 °C until use.

**FACS and NGS Analyses**

Cell characterization and sorting were performed using Fluorescence-Activated Cell Sorting (FACS, BD-Facsaria III), following the methodology previously described by Chao et al (2). The concentrations of Ras-GTP and Ras-GDP in each experiment were optimized to yield steep and comparable slopes of the binding signals, enabling clear separation of Nore1A mutants with different binding affinities in subsequent experiments. Preparations included Ras–GTP (GppNHp) and Ras–G12D–GTP (GppNHp) at final concentrations of 3 μM, and Ras–GDP and Ras–G12D–GDP at final concentrations of 15 μM and 25 μM, respectively. TBSA buffer (20 mM Tris-HCl, pH 7.5, 100 mM NaCl supplemented with 1% BSA) were used and tailored to the biochemical properties of the protein under analysis. This procedure enabled the selection of a WT gate defined to encompass approximately 99% of the Ras-binding clones in the sorting experiments with Nore1A$^{WT}$ (shown on Figure 1C and D and Supplementary Figure S1 A and C). Subsequently, upon application of the mutant library, four distinct cell populations were selected based on uniform expression levels but differing binding affinities. These populations were classified as high (HI), wild-type–like (WT), slightly lower (SL), and very low (LO) affinity, using the WT cell interaction as the reference. Cell sorting was performed using a 70-µm nozzle at 4 °C, with sample pressure set to 9, frequency at 87, and a flow rate threshold corresponding to approximately 8,000 events per second. Approximately 10 million cells were collected across all four sorted populations. The four sorted cell populations and 1 mL of unsorted control cells were separately inoculated into 100 mL of SD-CCA medium supplemented with kanamycin, chloramphenicol, ampicillin, and tetracycline, and grown at 30 °C with shaking at 225 rpm for 62 h. Plasmid DNA was extracted using the Zymoprep Yeast Plasmid Miniprep II Kit (Zymo Research). DNA concentrations were measured using a NanoDrop spectrophotometer (Thermo Scientific). DNA samples with high salt concentrations were treated with Pellet Paint® Co-Precipitant (Merck) to remove excess salt prior to sequencing process. For subsequent Next-Generation Sequencing (NGS) analysis, samples were prepared in duplicate from independent FACS-sorting experiments, ensuring data reproducibility and reliability. The extracted plasmid DNA was used as a template in PCR reactions to amplify the genetic material and to incorporate platform-specific adapters and promoter sequences compatible with Illumina sequencing technologies. Sequencing was carried out on Illumina MiSeq or NextSeq platforms, targeting a minimum read depth of $10^6$ unique reads per sample, processed in duplicate, to ensure analytical robustness and sufficient coverage for downstream analyses. Paired-end reads obtained from NGS were processed using the Galaxy web server (https://usegalaxy.org/). Sequence assembly and quality control were performed using the FLASH and FastQC tools, respectively. Quantitative analysis of the sequencing data, including the identification and enumeration of single and double mutants, was conducted using MATLAB (MathWorks). Further data analysis was conducted using Microsoft Excel and custom scripts written in Python (version 3.8), executed in the Jupyter

Notebook environment (version 6.1.4). Graphical representations and statistical analyses were performed using OriginPro software (Version 2023, OriginLab Corporation, USA).

**NGS data analysis**

Paired-end reads obtained from NGS experiments were merged, and sequence quality was evaluated using the FastQC tool (https://www.bioinformatics.babraham.ac.uk/projects/fastqc/). Subsequent analysis was conducted using a custom MATLAB script developed to identify single and double mutation events. For single mutation analysis, reads were aligned to the WT Nore1A sequence, and those containing more than two mutations were discarded. In addition, we discarded the sequences that occurred less than 10 times in the naïve library to increase the accuracy of our analysis. For each Nore1A variant sequence, the number of occurrences was counted in both the sorted and the naïve populations, and its frequency ($f^i$) in the particular gate was calculated (Eq. 1).

$$f^i = \frac{count^i}{count_{total}} \quad (1)$$

Using the relative frequencies in a sorted population and the naïve library, the enrichment ($e^i$) of each Nore1A mutant was then computed (Eq. 2):

$$en^i = \ln(\frac{(f^i)_{sorted}}{(f^i)_{naïve}}) \quad (2)$$

The enrichment values were further normalized in each gate to range from -1 to +1 using the maximum and the minimum enrichment values in this gate $en^{max}$ and $en^{min}$, respectively (eq. 3):

$$en^i_{norm} = 2\frac{en^i - en^{min}}{en^{max} - en^{min}} - 1 \quad (3)$$

We furthermore, excluded the Nore1A variants for which enrichment values were available for less than 3 gates. When only one enrichment value was missing, it was imputed using the CatBoost algorithm based on the corresponding values of the three variants with the most similar enrichment profiles.

To predict $\Delta\Delta G_{bind}$ for all detected Nore1A variants from enrichment values in four gates ($X_1$, $X_2$, $X_3$, $X_4$), we utilized a non-linear machine learning approach based on Extremely Randomized Trees (Extra-Trees)(3). The model was trained on 38 experimental $\Delta\Delta G_{bind}$ datapoints summarized in Supplementary Table 1. To account for the four binding states represented in the dataset, we implemented a state aware ensemble in which mutations were first assigned to their dominant affinity class HI ($X_1$), WT ($X_2$), SL ($X_3$), or LO ($X_4$) and subsequently evaluated using a corresponding set of specialized Extra-Trees regressors(4). This formulation allows each class of mutations to be modelled within its appropriate energetic context, preventing the model from averaging across very different mutation types and underestimating strongly stabilizing or destabilizing effect. Feature importance scores were computed within each state specific model to quantify the contribution of each enrichment feature to $\Delta\Delta G_{bind}$. The trained model was tested in the leave-one-out cross validation setting on our experimental dataset. The trained model was then applied to predict $\Delta\Delta G_{bind}$ values for all Nore1A single and double mutants detected by NGS when interacting with RasWT-GTP, RasG12D-GTP, RasWT-GDP and RasG12D-GDP.

The coupling energy $\Delta G_i$ upon two mutations X and Y was calculated according to:

$$\Delta G_i = \Delta\Delta G_{bind}^{X} + \Delta\Delta G_{bind}^{Y} - \Delta\Delta G_{bind}^{XY} \qquad (4)$$

Here, $\Delta\Delta G_{bind}^{X}$ and $\Delta\Delta G_{bind}^{Y}$ represent the change of the binding free energy of the single mutants X and Y, respectively, $\Delta\Delta G_{bind}^{XY}$ represents the change of binding free energy of the double mutant containing mutations X and Y. The mutation was defined as exhibiting negative epistasis if $\Delta G_i$ was negative within the uncertainty of the predictions, that is:

$$\Delta G_i = \Delta\Delta G_{bind}^{X} + \Delta\Delta G_{bind}^{Y} + \sqrt{\delta\Delta\Delta G_{bind}^{X}{}^{2} + \delta\Delta\Delta G_{bind}^{Y}{}^{2}} - \Delta\Delta G_{bind}^{XY} + \delta\Delta\Delta G_{bind}^{XY} < 0$$

(5)

Here, $\delta\Delta\Delta G_{bind}^{X}$, $\delta\Delta\Delta G_{bind}^{Y}$ and $\delta\Delta\Delta G_{bind}^{XY}$ are uncertainties in prediction of $\Delta\Delta G_{bind}$ for mutation X, Y, and XY, respectively.

The mutation was defined as exhibiting positive epistasis if $\Delta G_i$ was positive within the uncertainty of the predictions, that is:

$$\Delta G_i = \Delta\Delta G_{bind}^{X} + \Delta\Delta G_{bind}^{Y} - \sqrt{\delta\Delta\Delta G_{bind}^{X}{}^{2} + \delta\Delta\Delta G_{bind}^{Y}{}^{2}} - \Delta\Delta G_{bind}^{XY} - \delta\Delta\Delta G_{bind}^{XY} > 0$$

(6)

**NORE1A Mutant Construction to $K_D$ evaluation**

Gene constructs encoding NORE1A mutants (Supplementary Table 1) were generated from the wild-type (WT) NORE1A sequence using the Transfer-PCR (TPCR) methodology (5). Nore1A WT and mutants were expressed in BL21(DE3) *E. coli* at 15 °C overnight, purified on a Ni beads column (Adar Biotech, Rehovot, Israel). Protein purity was verified by gel electrophoresis (Figure S3) and the concentration was measured by absorbance at 280 nm and the Bradford assay. Protein samples were aliquoted and stored at -80°C.

**GDI Assay to Evaluate the Effect of Nore1A on Ras Nucleotide Dissociation**

To measure Kd of Nore1A mutants to RasWT-GTP and RasG12D-GTP, a guanine nucleotide dissociation inhibition (GDI) assay was performed using fluorescently labeled mant-nucleotides (mant-GDP or mant-GppNHp) (Jena Bioscience). Initially, purified Ras protein was buffer-exchanged to reduce the concentration of $MgCl_2$, which is required to destabilize nucleotide binding and enable exchange. This was performed using a centrifugal (Amicon) concentrator with a low-$Mg^{2+}$ buffer (20 mM Tris-HCl pH 7.5, 50 mM NaCl, 1 mM DTT, 0.1 mM $MgCl_2$). For nucleotide exchange (6), Ras was incubated with a 20-fold molar excess of mant-GDP or mant-GppNHp at 4°C for 60 minutes. The exchange reaction was terminated by the addition of $MgCl_2$ to a final concentration of 10 mM to reestablish Ras–nucleotide complex stability ) at 4°C for 30 minutes (6). Excess unbound nucleotide was removed by passing the sample through a Zeba™ Spin Desalting Column (Thermo Fisher Scientific), previously equilibrated with the assay buffer (20 mM Tris-HCl pH 7.5, 100 mM NaCl, 2 mM DTE, 5 mM $MgCl_2$). Recombinant Nore1A was prepared in the same assay buffer and diluted to a range of final concentrations (e.g., 0.025–10 μM) to assess its effect on Ras nucleotide dissociation. The Ras-mant-nucleotide complexes were then incubated without Nore1A variant or with Nore1A at a variable concentration at 37 °C for 5 minutes. To initiate the nucleotide exchange reaction, a 1000-fold molar excess of unlabeled nucleotide (GDP or GppNHp) (Abcam) was added to each sample. The dissociation of the mant-labeled nucleotide from Ras over time was monitored by measuring fluorescence, using excitation at 355 nm and emission at 448 nm. Each experiment performed with a certain Nore1A variant concentration was analyzed first to determine the observed time constant τ by fitting an exponential decay function to the fluorescence intensity as a function of time. $k_{obs}$ was then calculated as 1/τ for each experiment. Next, $k_{obs}$ was plotted vs. the Nore1A variant concentration and fitted to a quadratic equation as described in a previous study:

$$k_{\{obs\}} = k_{\{-1\}} - \left(k_{\{-1\}} - k_{\{-2\}}\right) * \frac{([R_0]+[E_0]+K_d) - \sqrt{\{([R_0]+[E_0]+K_d)^2 - 4*[R_0]*[E_0]\}}}{2*[R_0]} \quad (7)$$

where $k_{\{-1\}}$ and $k_{\{-2\}}$ are the nucleotide dissociation rate constants from free Ras and from the Ras–Nore1A complex, respectively, $K_D$ is the binding affinity fitted in the experiment, and $[R_0]$ and $[E_0]$ denote total concentrations of Ras and Nore1A variant, respectively. $[R_0]$ of 0.2 μM was used in all experiments. $K_D$ values were converted into $\Delta\Delta G_{bind}$ according to the formula:

$$\Delta\Delta G_{bind} = RTLn\left(\frac{K_{D\_WT}}{K_{D\_MUT}}\right) \quad (8)$$

Additional data for $\Delta\Delta G_{bind}$ was taken from another study that measured $K_D$s of Nore1A mutants to RasWT-GTP using the same GDI assay (7). To adjust for different ionic strength conditions in the previous study, we multiplied all $\Delta\Delta G_{bind}$ values by 0.6.

**Molecular Dynamics Simulation Protocol**

All molecular dynamics (MD) trajectories were generated utilizing the OpenMM software suite [38]. Initial structural coordinates for the complex were obtained from the Protein Data Bank (PDB ID: 3DDC). To prevent the large movements in a flexible loop of Nore1A (residues 244–278) that does not participate in Ras binding and is unresolved in available crystal structures, we truncated the loop to 8 residues (244–248 and 274–276) by connecting P248 to S274. We predicted the conformation of Nore1A with the truncated loop using AlphaFold. A guanosine triphosphate (GTP) molecule and its coordinating magnesium ion ($Mg^{2+}$) were docked into the binding pocket, adopting the spatial configuration and binding coordinates detailed in(8). System parameterization was conducted with the AMBER14 force field, utilizing customized small-molecule parameters for the GTP–$Mg^{2+}$ cofactor. The complex was explicitly solvated with the rigid TIP3P-FB water model. To compute long-range electrostatic interactions under periodic boundary conditions, the Particle Mesh Ewald (PME) algorithm was applied, using an Ewald error tolerance of 0.0005 and a 1.0 nm real-space cutoff. A 2.0 fs integration timestep was enabled by constraining the lengths of all hydrogen-involving bonds. Simulations were run in the isothermal-isobaric (NPT) ensemble. A target temperature of 300 K was maintained using a LangevinMiddleIntegrator with a friction coefficient of 1.0 $ps^{-1}$ to provide precise kinetic sampling. The protocol was initiated with a rigorous energy minimization phase to eliminate steric clashes and structurally optimize the $Mg^{2+}$ coordination sphere alongside the

GTP phosphate backbone. Atomic velocities were then assigned from a 300 K Maxwell-Boltzmann distribution, and the system underwent NPT equilibration to ensure full structural relaxation. Following equilibration, a 50 ns production simulation was executed, corresponding to $2.5 \times 10^7$ integration steps. Trajectory frames were written at regular intervals for downstream analysis. System stability and equilibrium throughout the 50 ns sampling period were validated by continuously monitoring the root-mean-square deviation (RMSD), as well as thermodynamic fluctuations in temperature, potential energy, and overall system density. After the simulation, the trajectory file was analysed by calculating root-mean-square fluctuation (RMSF) of Cα atoms to assess local residue flexibility over 50 ns simulations for all the residues in the Ras-Nore1A complex.

#### Supplementary Figures and Legends

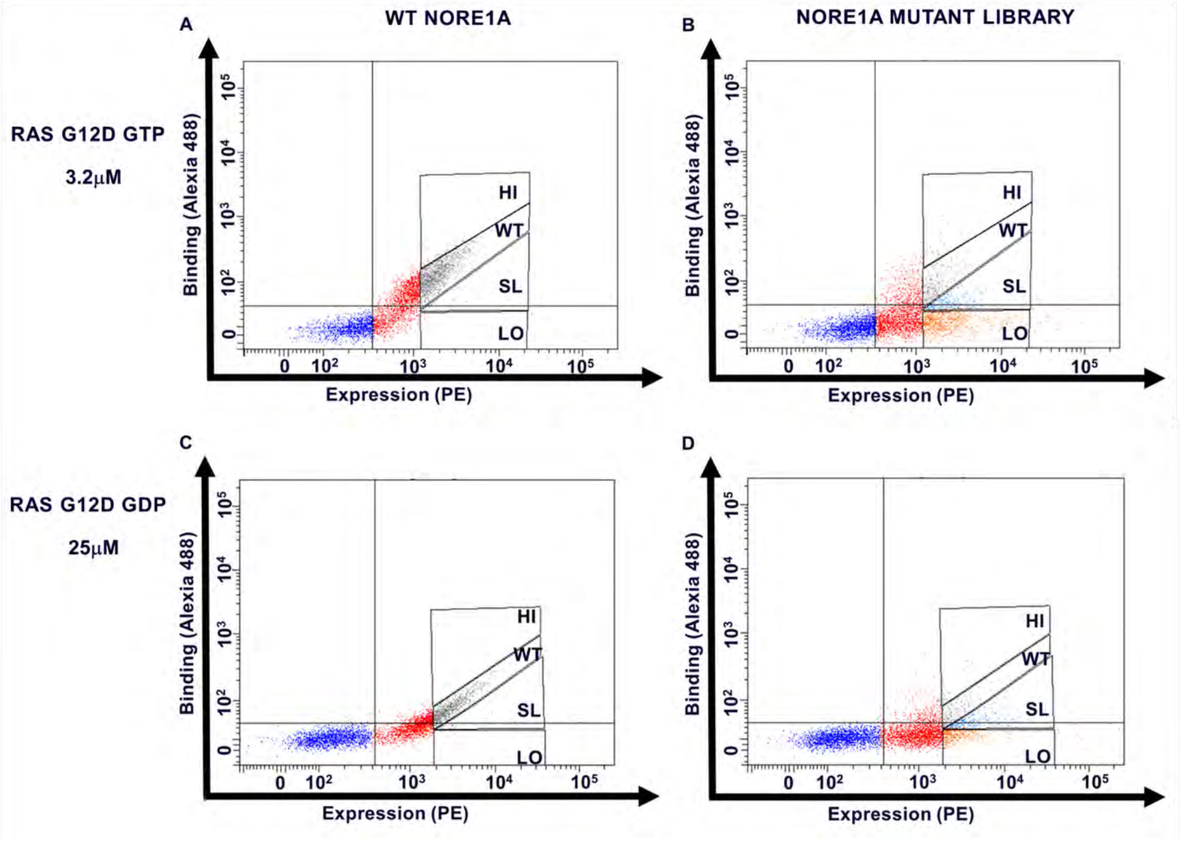


**Figure S1:** FACS data showing binding of Nore1A WT or Nore1A library to Ras. The PE signal monitoring Nore1A expression is shown on the X-axis and the Alexa signal monitoring binding between fluorescently-labelled Ras and Nore1A variants expressed on the yeast surface is shown on the y-axis. (A) Nore1A WT binding to Ras-G12D-GTP (3.2 μM). (B) Nore1A library binding to Ras-G12D-GTP (3.2 μM). (C) Nore1A WT binding to RasG12D-GDP (25 μM). (D) Nore1A library binding to RasG12D-GDP (25 μM). Four sorting gates are depicted: HI (with affinity higher than WT), WT (with wild-type like affinity), SL (with slightly lower than wild-type affinity), LO (with much lower than wild-type affinity).

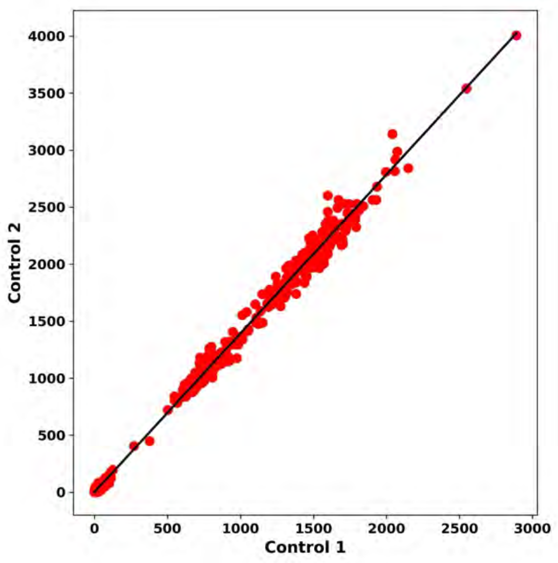


**Figure S2:** Correlation between counts of sequences detected by NGS for two identical samples corresponding to the naïve library containing single and double mutations. R-value between the two samples is calculated to be 0.98.

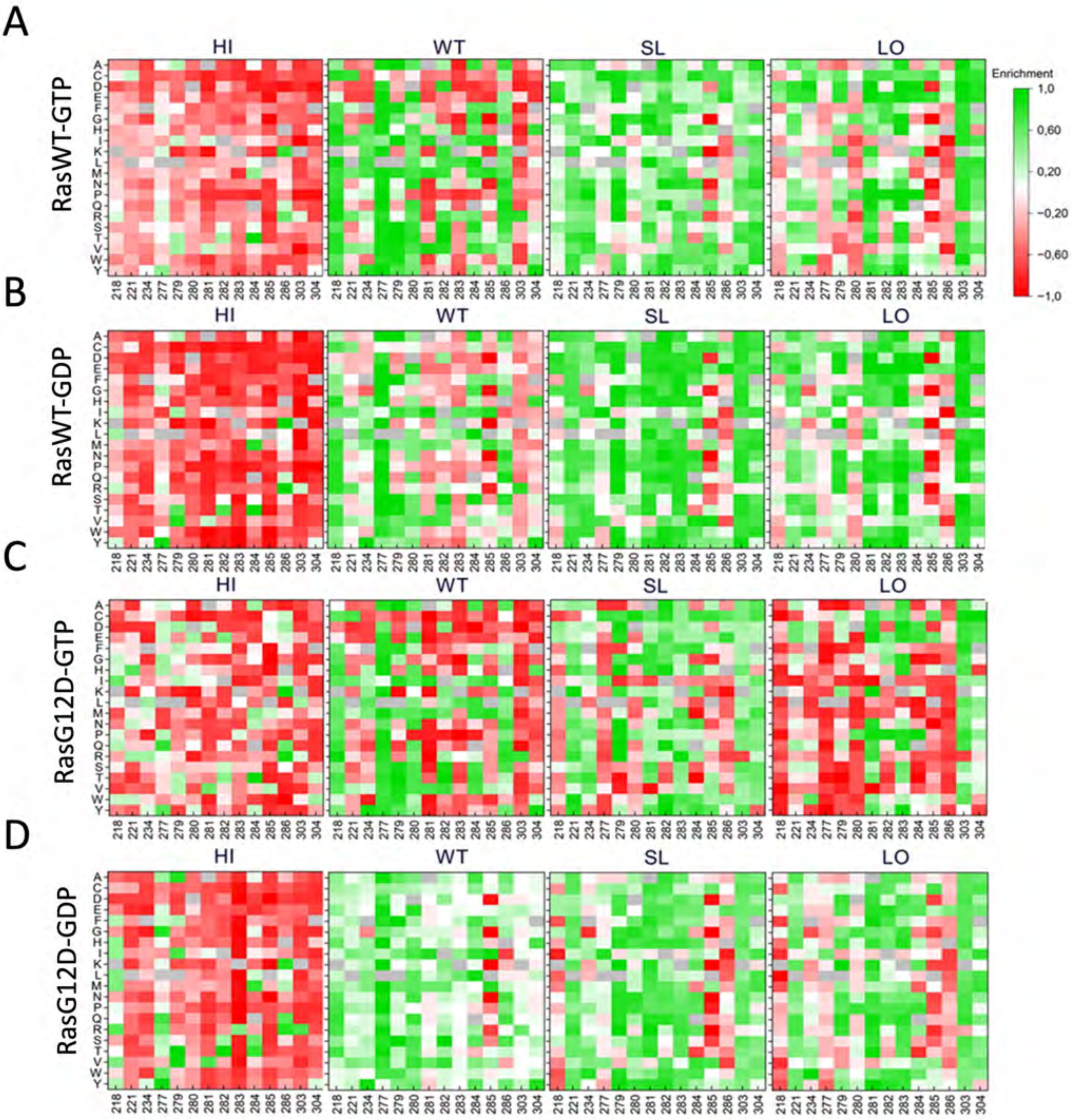


**Figure S3:** Heatmaps showing normalized enrichment values for Nore1A single mutants binding to four Ras variants: RasWT-GTP (A), RasWT-GDP (B), RasG12D-GTP (C), RasG12D-GDP (D). Here, twenty amino acids are depicted on the Y acid and randomized positions on Nore1A – on the X axis. The heatmaps show normalized enrichment of each variant in the high-affinity (HI) gate, wild-type like affinity (WT), slightly lower affinity (SL) and very low affinity (LO) varying from +1 (green) to -1 (red).

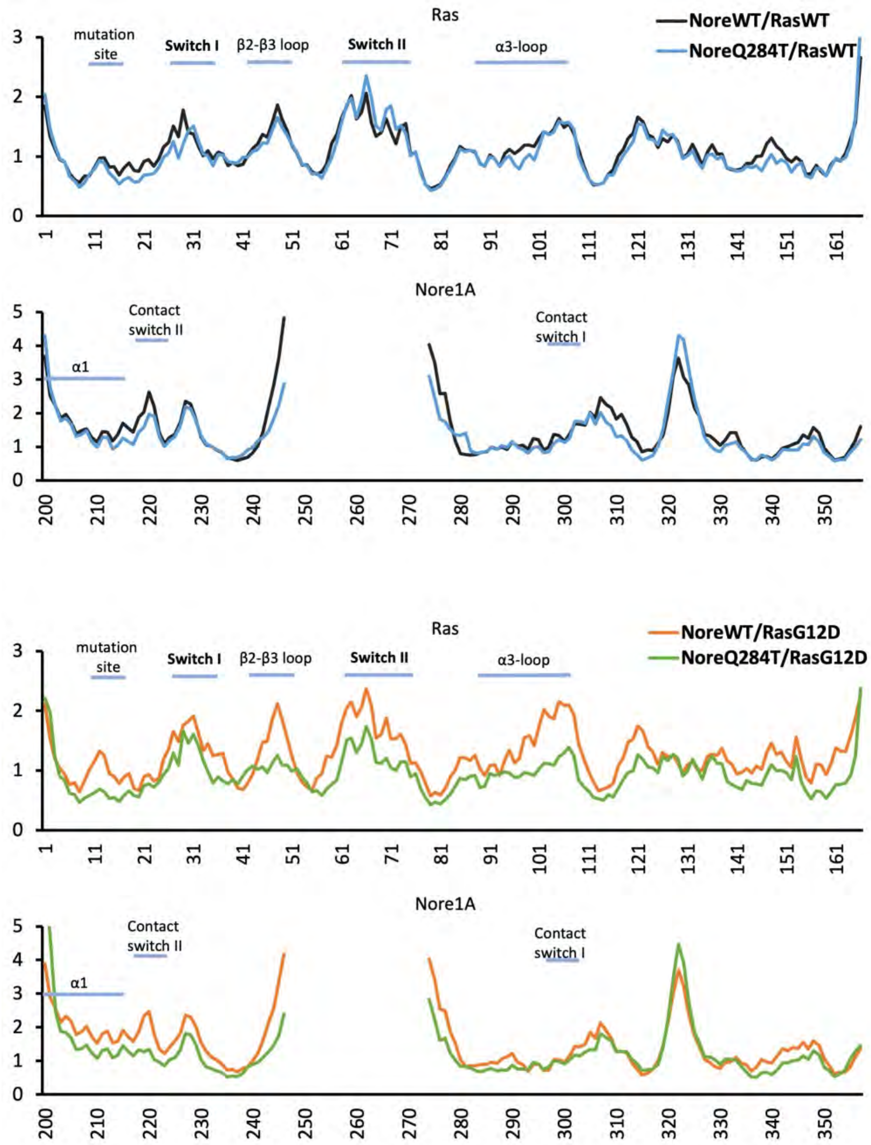


**Figure S4:** RMSF of Ras/Nore1A variant complexes. pairwise comparison of two mutant complexes: (A) NoreWT and NoreQ284T interacting with RasWT; (B) NoreWT and NoreQ284T interacting with RasG12D. RMSF was calculated from 50 nsec MD simulation of the complexes done under identical conditions. Important regions on Ras and Nore1A are marked about the RMSF curves.

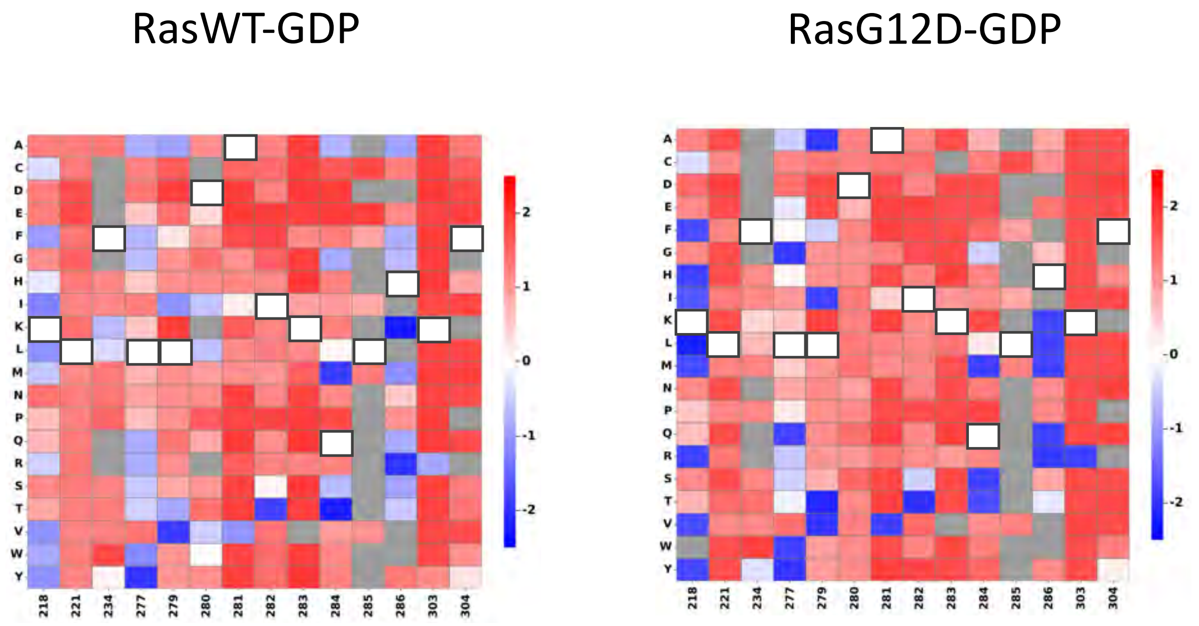


**Figure S5:** Quantitative binding landscapes for all single mutations for Nore1A-RasWT-GDP complex (C) and Nore1A/RasG12D-GDP complex (D). Y axis shows amino acid identity while X-axis shows position on Nore1A. Mutations are color coded from highly stabilizing (blue) to highly destabilizing (Red). In white squares are WT amino acids. In grey -- and mutations for which no predictions are available.

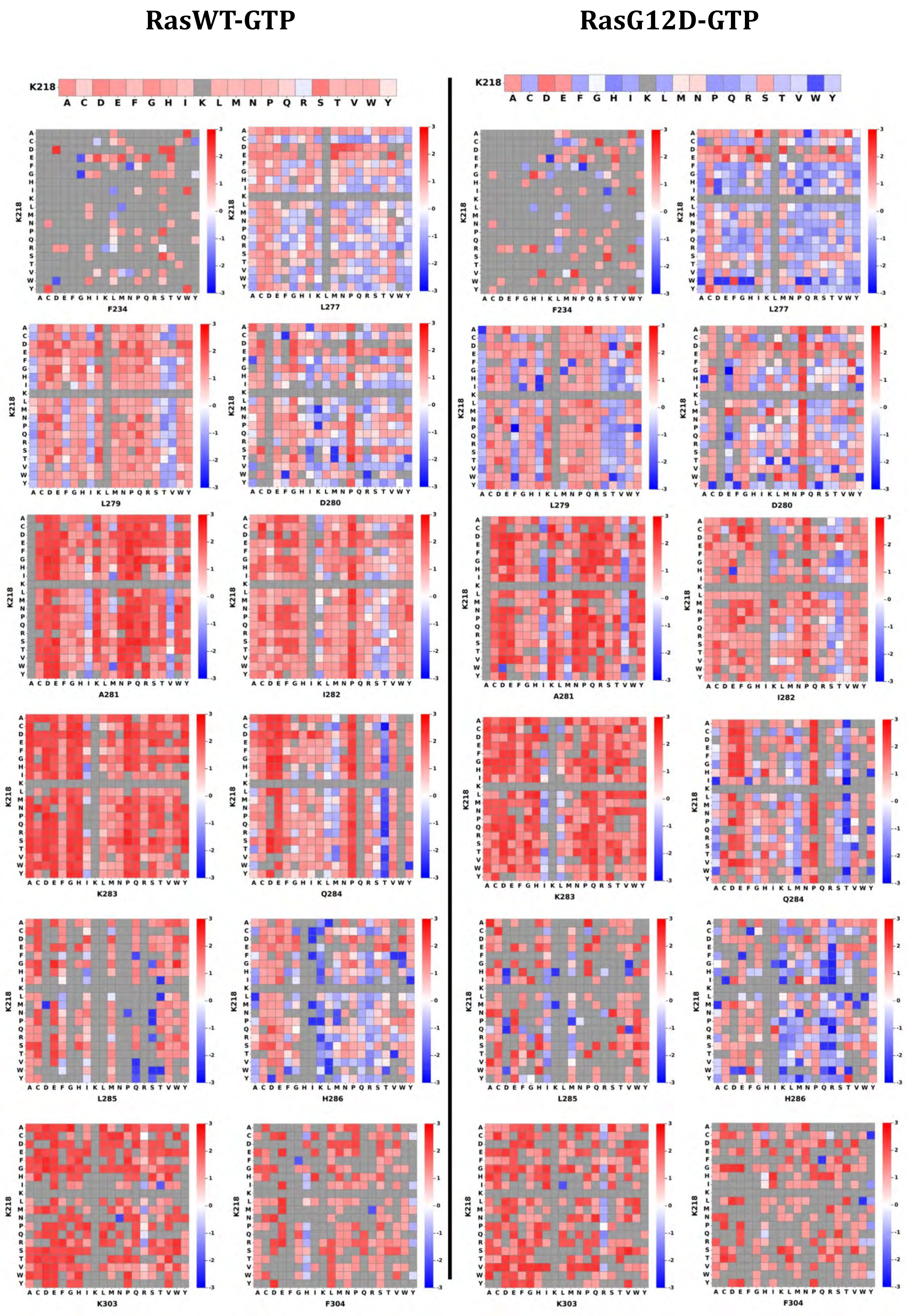

RasWT-GTP
RasG12D-GTP
K218
F234
L277
L279
D280
A281
I282
K283
Q284
L285
H286
K303
F304

### RasWT-GTP

### RasG12D-GTP

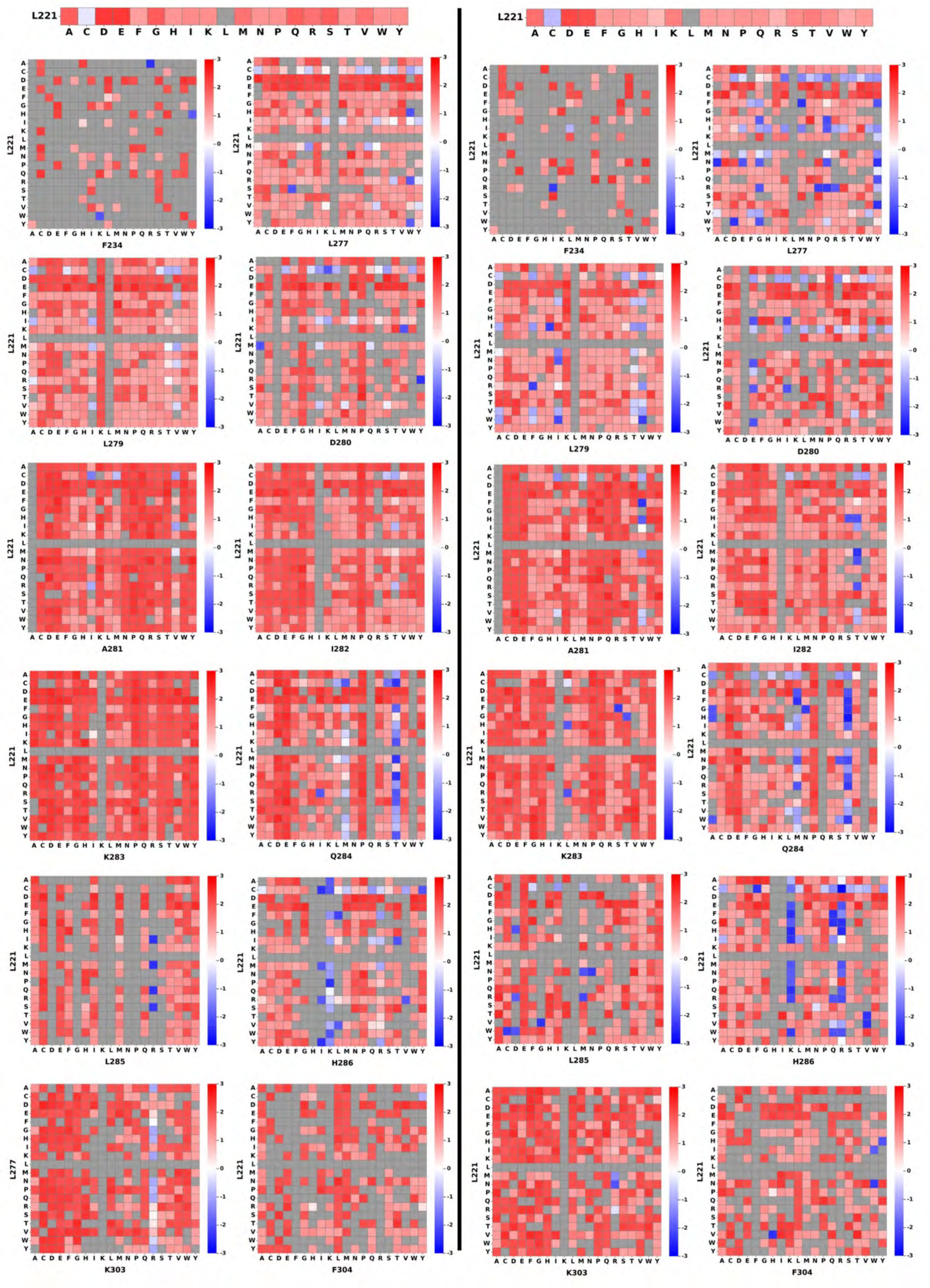

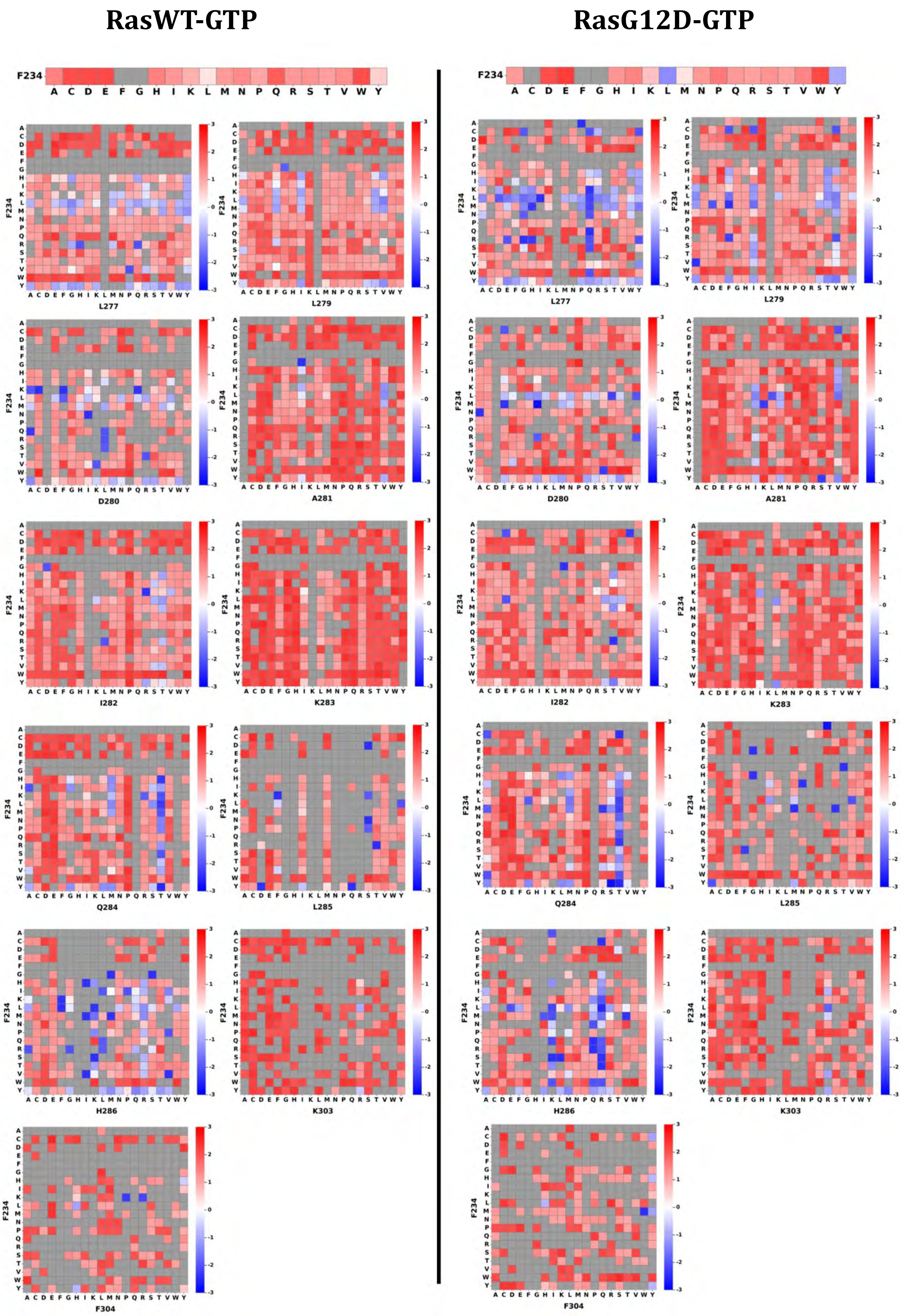
RasWT-GTP
RasG12D-GTP
F234
A C D E F G H I K L M N P Q R S T V W Y
L277
L279
D280
A281
I282
K283
Q284
L285
H286
K303
F304

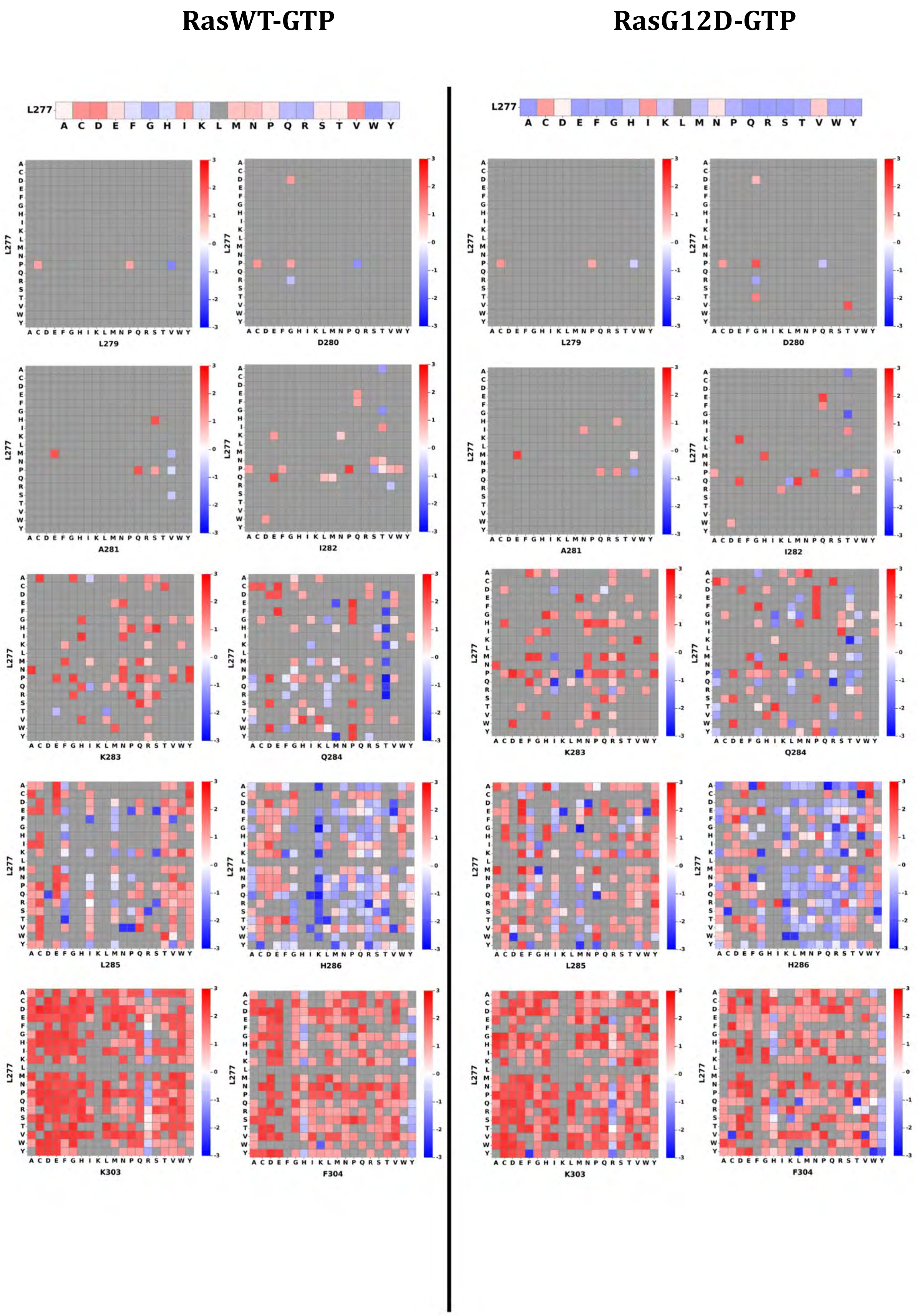

RasWT-GTP
RasG12D-GTP
L277
A C D E F G H I K L M N P Q R S T V W Y
L279
D280
A281
I282
K283
Q284
L285
H286
K303
F304

**RasWT-GTP**

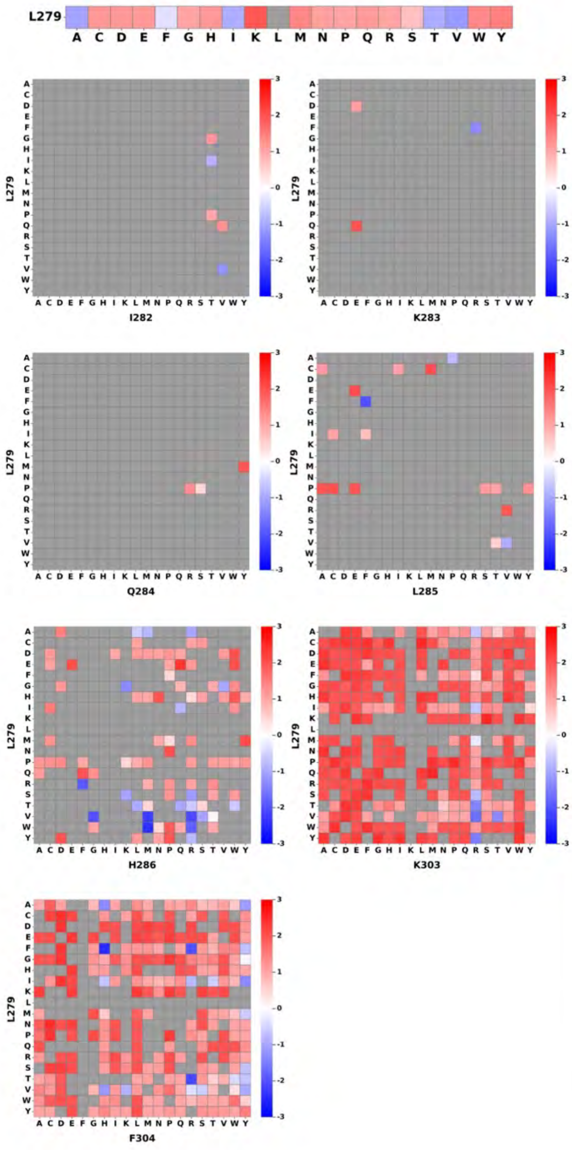


**RasG12D-GTP**

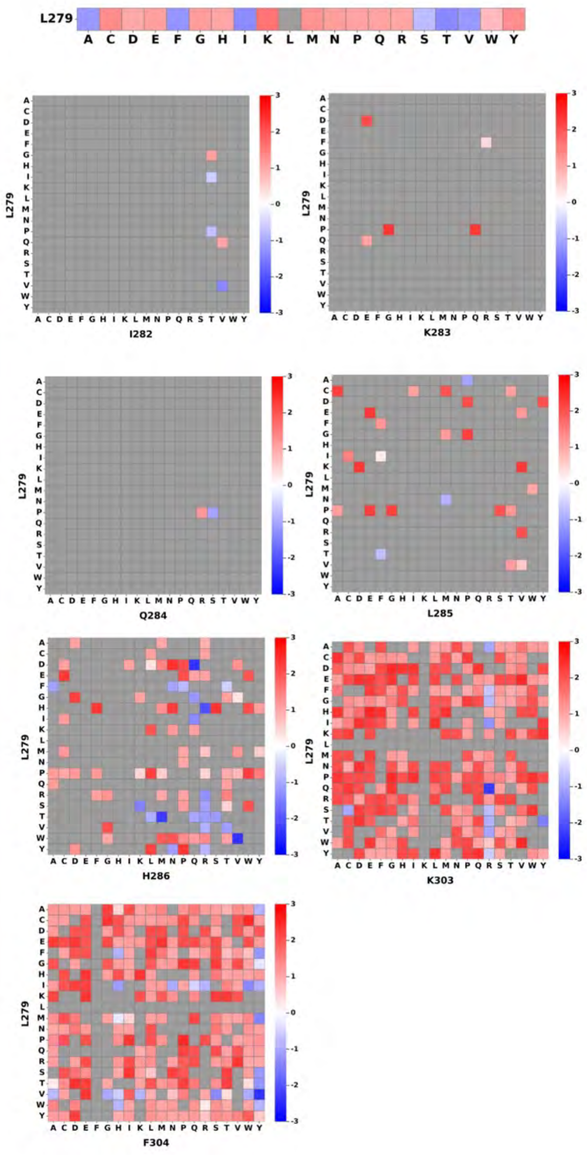

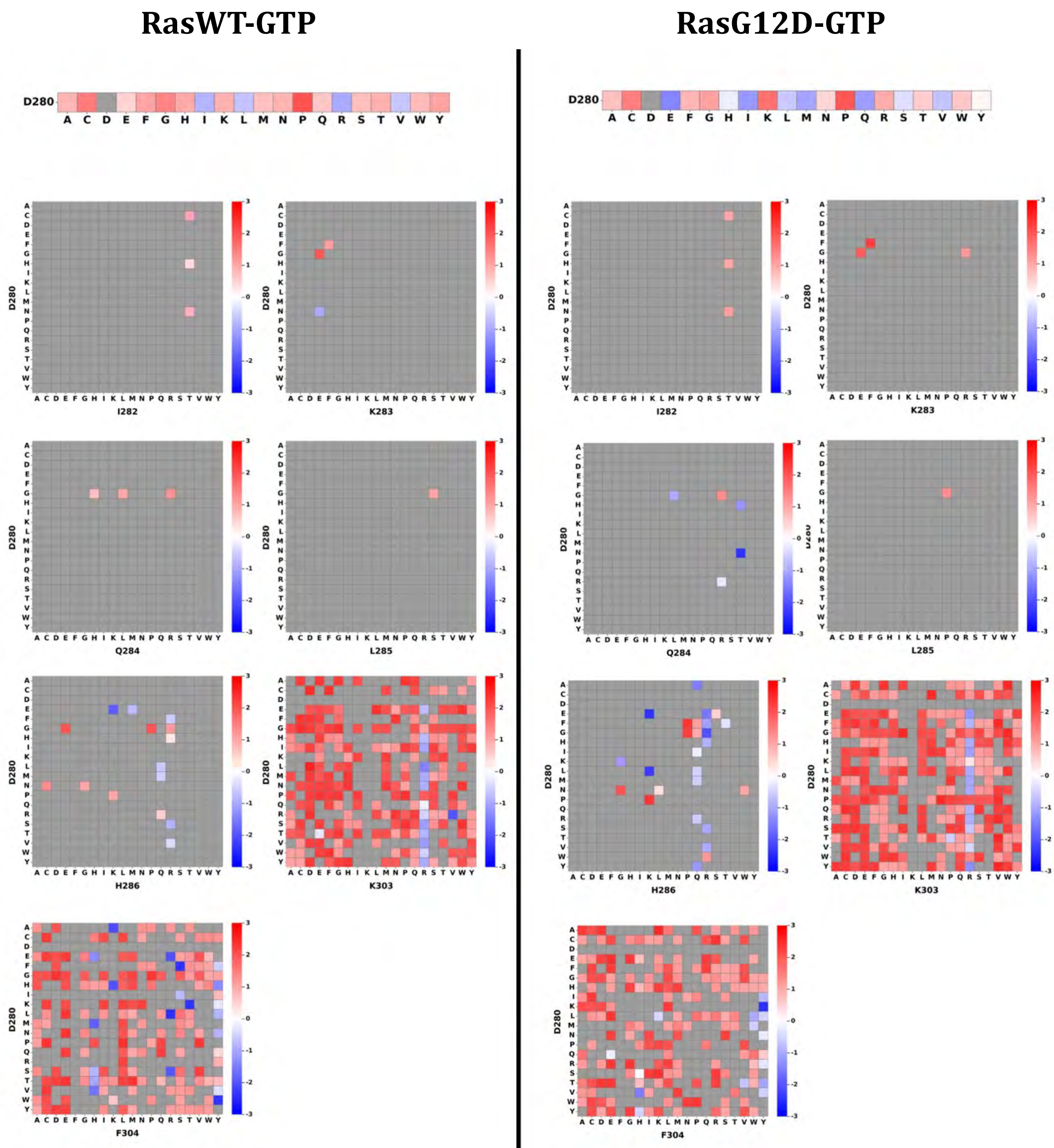
RasWT-GTP
RasG12D-GTP
D280
A C D E F G H I K L M N P Q R S T V W Y
I282
K283
Q284
L285
H286
K303
F304

#### RasWT-GTP

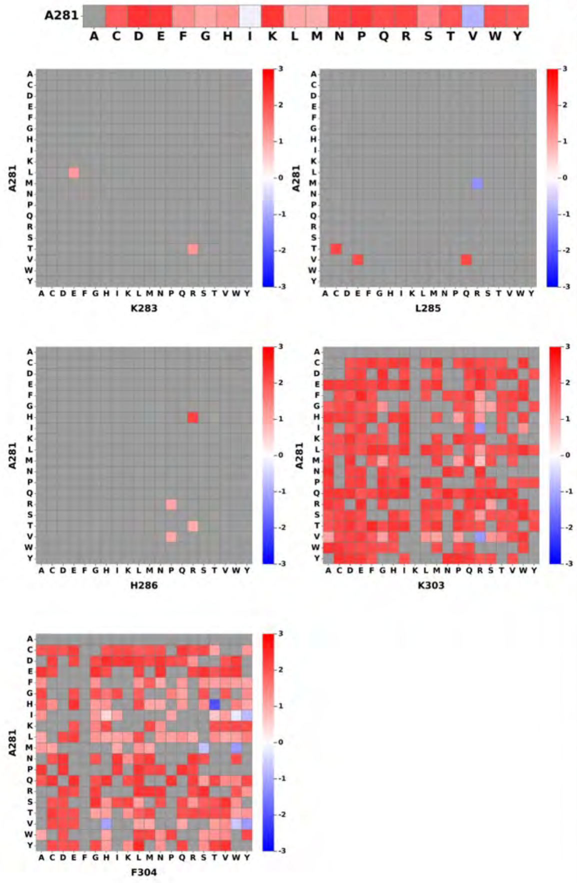


#### RasG12D-GTP

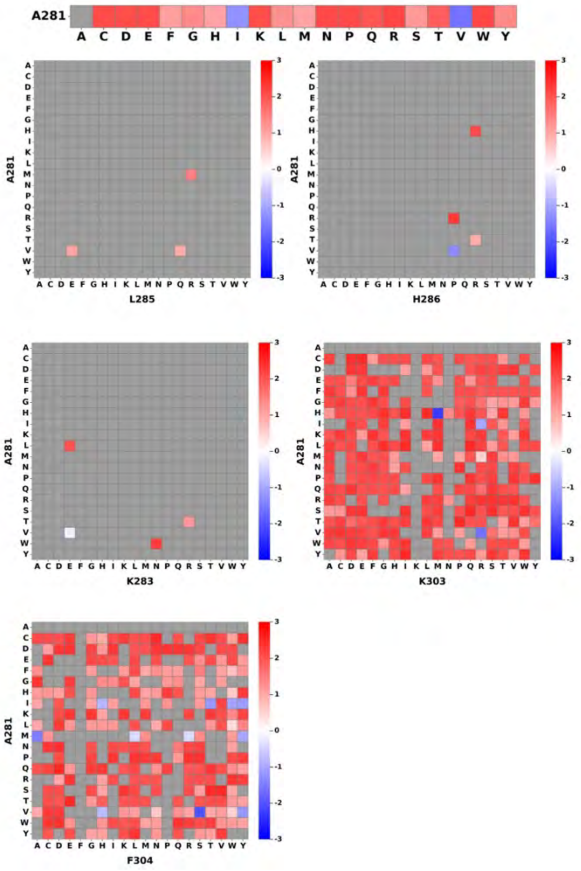

**RasWT-GTP** **RasG12D-GTP**

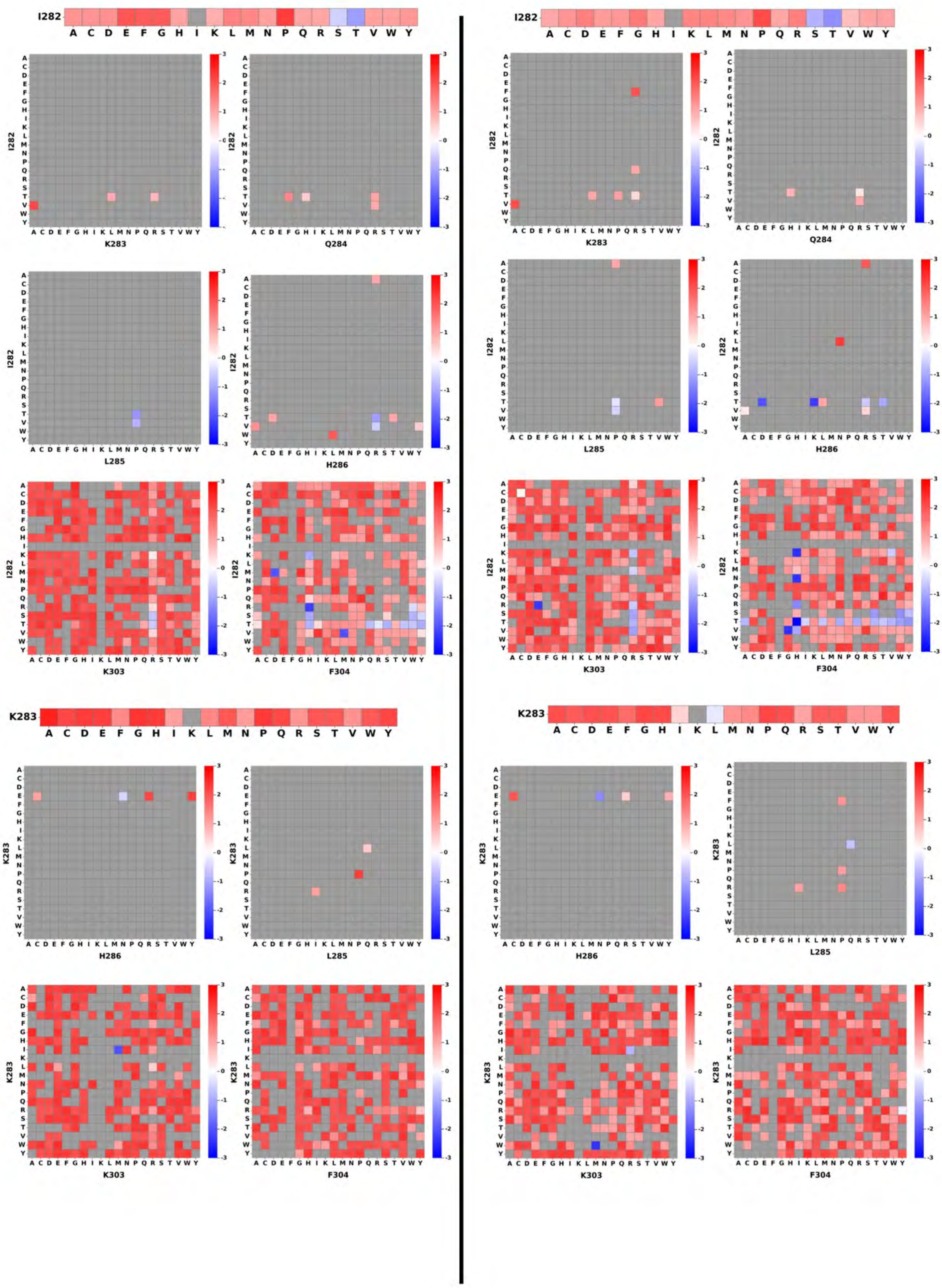

**RasWT-GTP** **RasG12D-GTP**

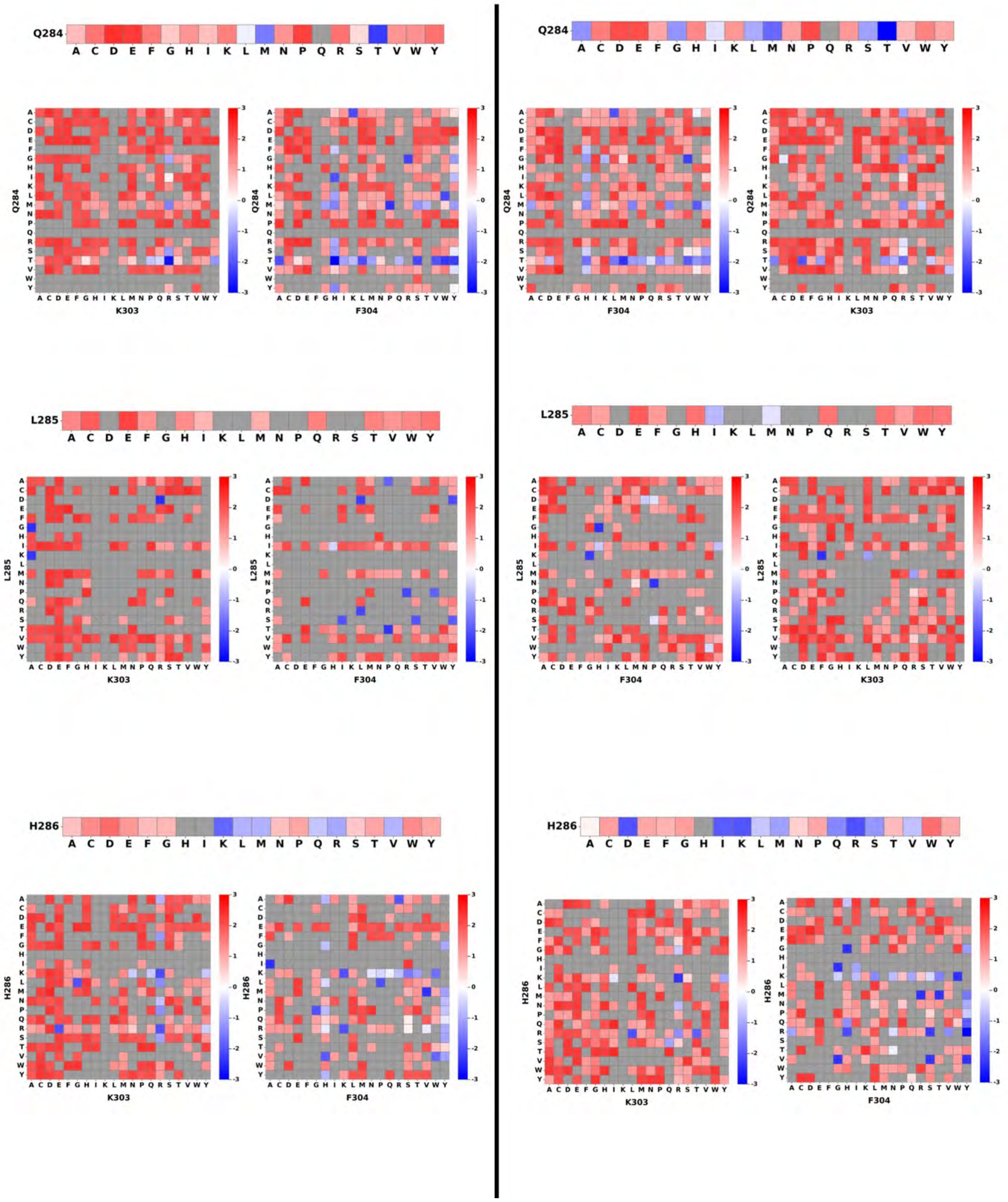


**Figure S6:** $\Delta\Delta G_{bind}$ values for all double mutations detected in the RasWT-GTP (left) and RasG12D-GTP (right) interactions. The first mutations is given on the Y axis and the second on the X axis. The $\Delta\Delta G_{bind}$ values are colored according to the legend from highly destabilizing in red to highly stabilizing in blue. The $\Delta\Delta G_{bind}$ values for the corresponding first single mutation is given on top for comparison.

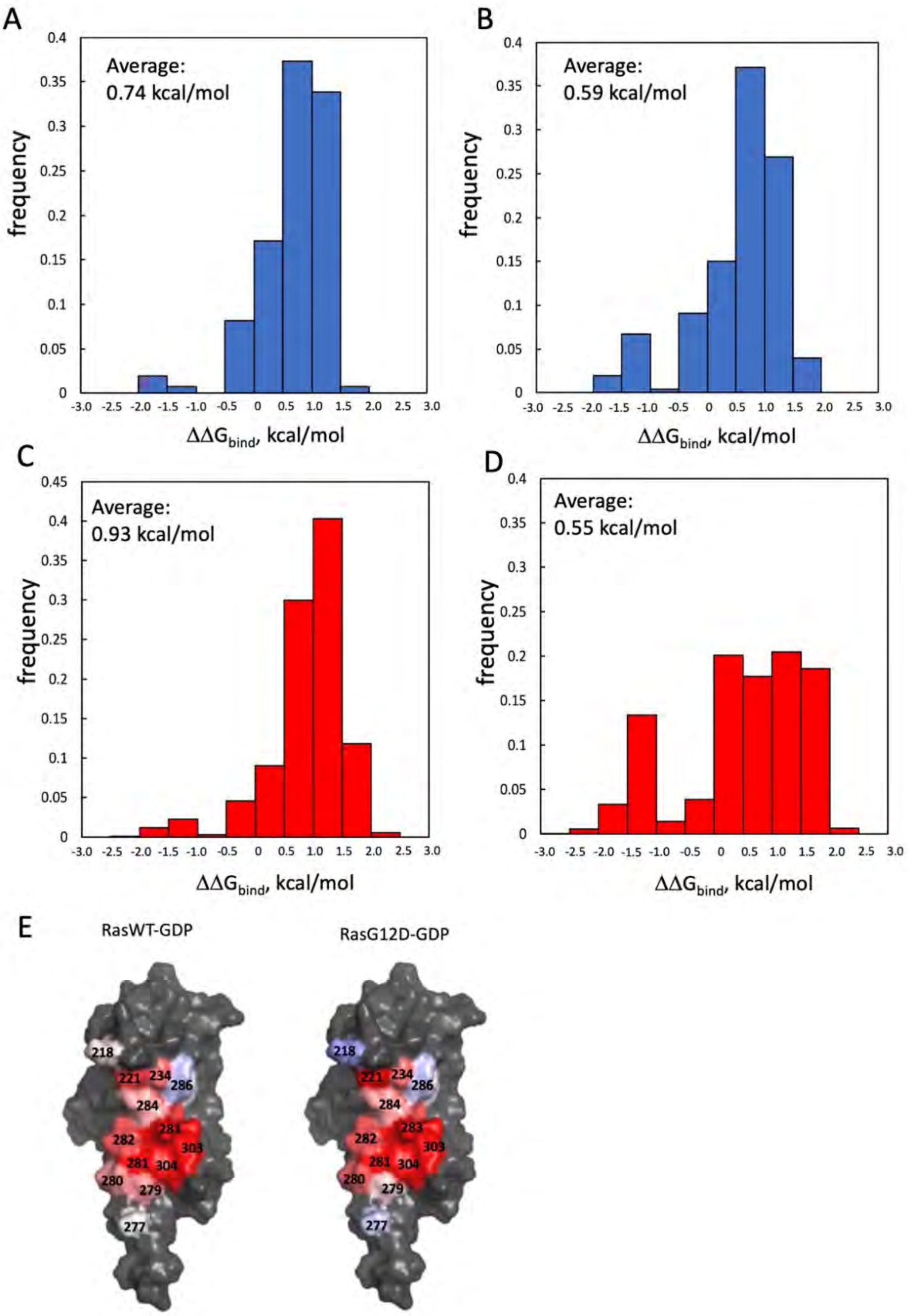


**Figure S7:** (A)-(D) Histograms of the predicted $\Delta\Delta G_{bind}$ values for all detected single and double mutations. (A) Single mutations for the Nore1A-RasWT-GTP interaction; (B) Single mutations for the Nore1A-RasG12D-GTP interactions; (C) Double mutations for the Nore1A-RasWT-GTP interaction; (D) Double mutations for the Nore1A-RasG12D-GTP interaction. (E) Average per position $\Delta\Delta G_{bind}$ value mapped on the structure of Nore1A interacting with RasWT-GTP (left) and RasG12D-GTP (right). The values vary from -1.0 kcal/mol (blue) to 1.0 kcal/mol (red).

**RasWT-GTP** **RasG12D-GTP**

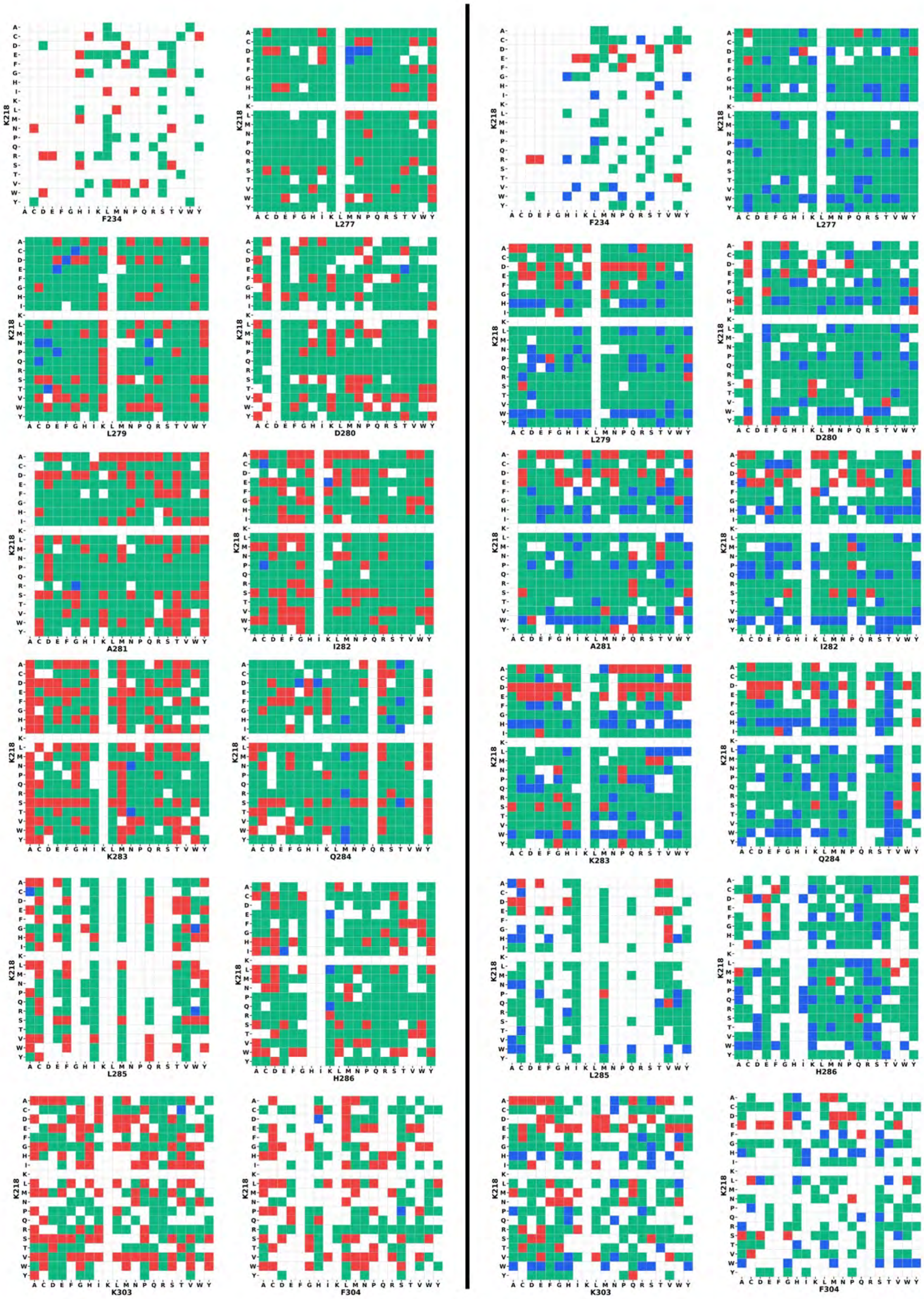

**RasWT-GTP** **RasG12D-GTP**

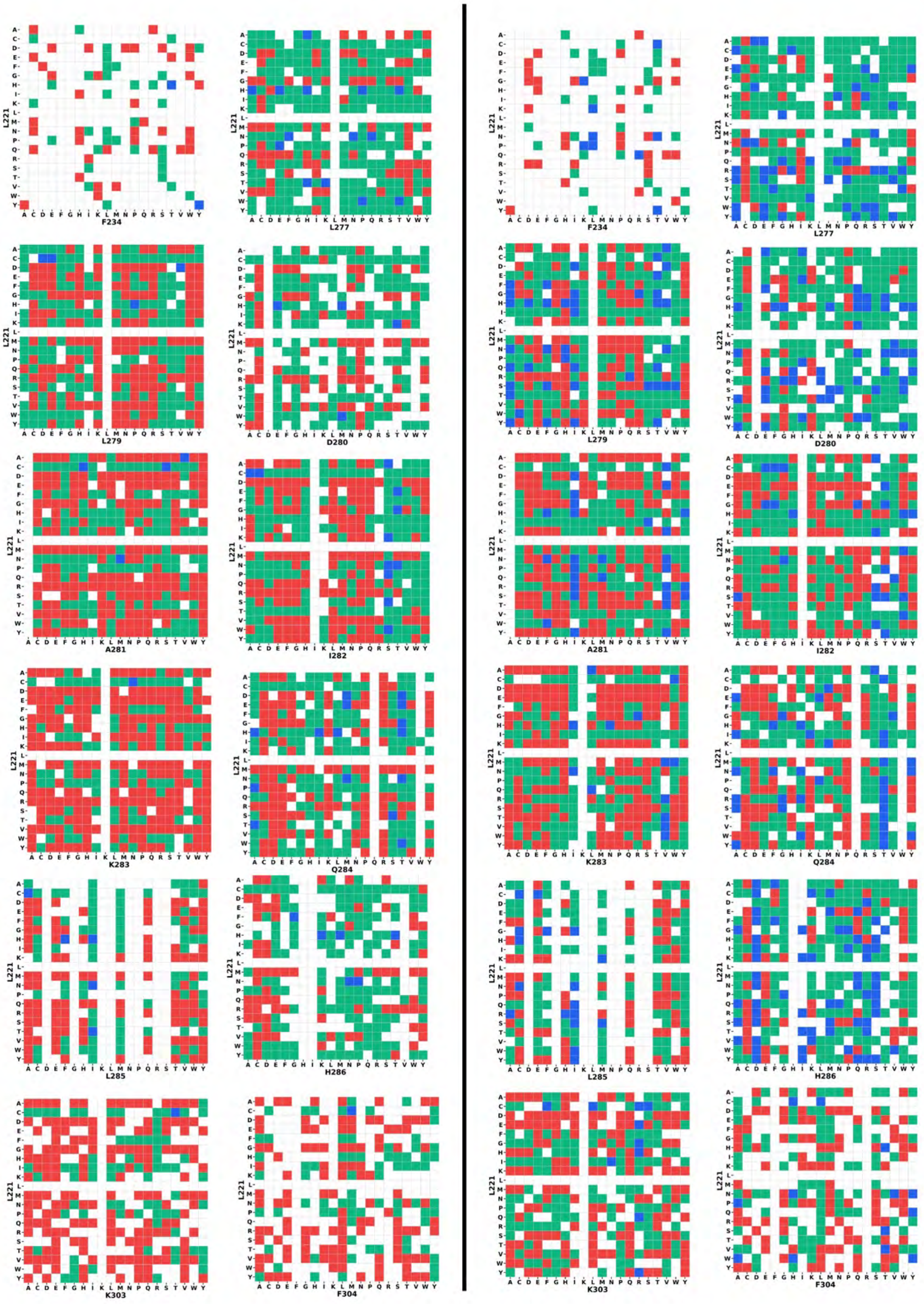

**RasWT-GTP** **RasG12D-GTP**

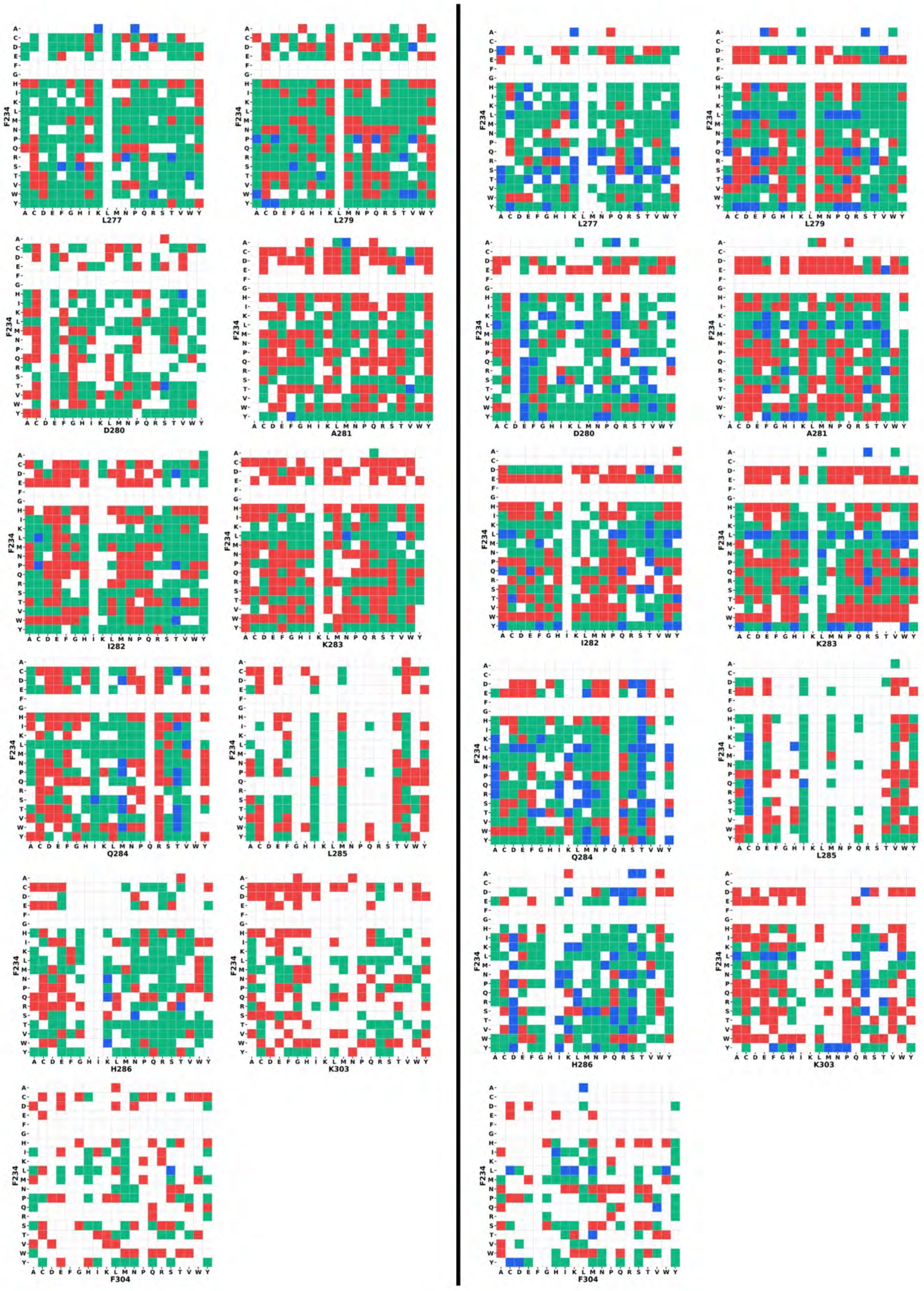

**RasWT-GTP**

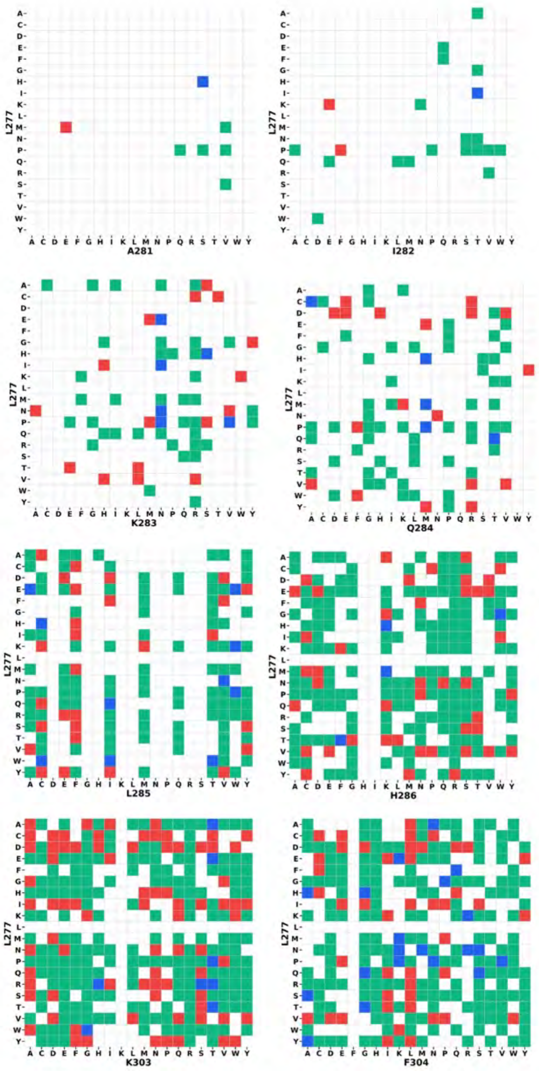


**RasG12D-GTP**

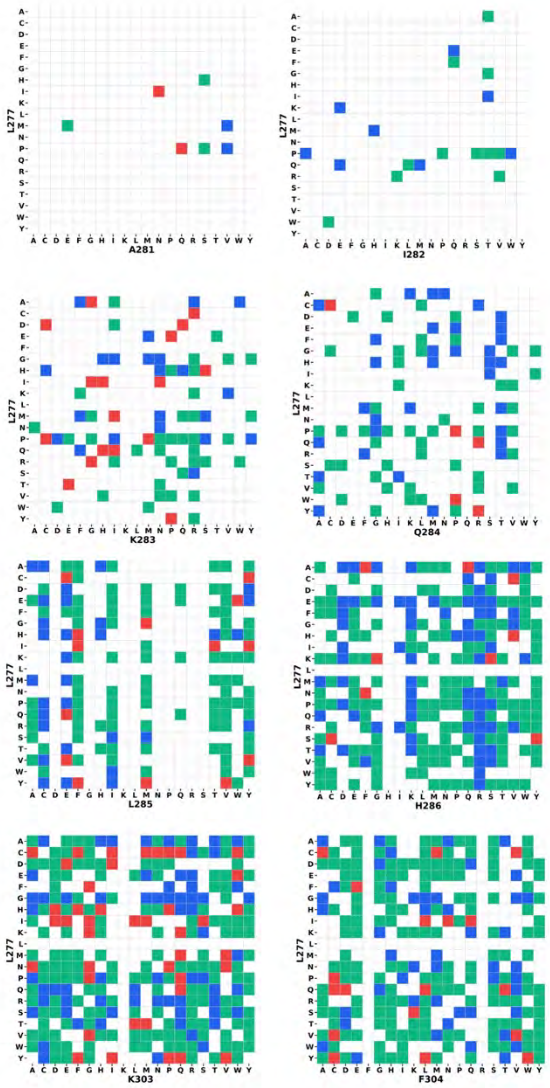

**RasWT-GTP**

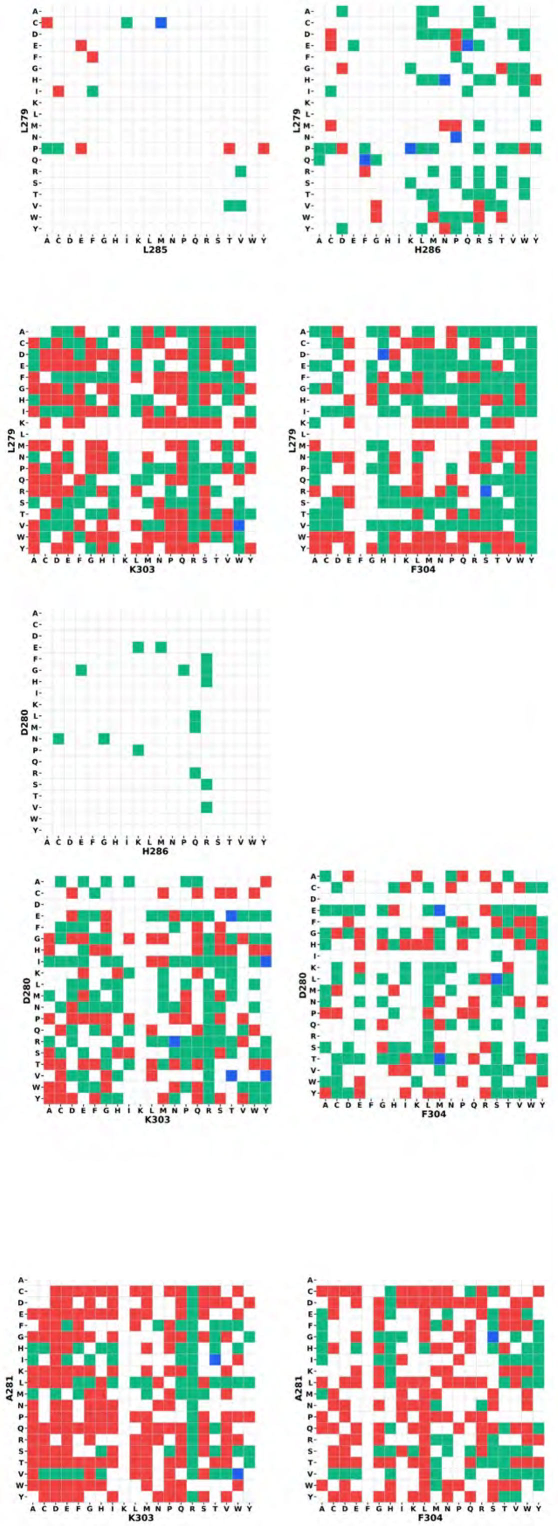


**RasG12D-GTP**

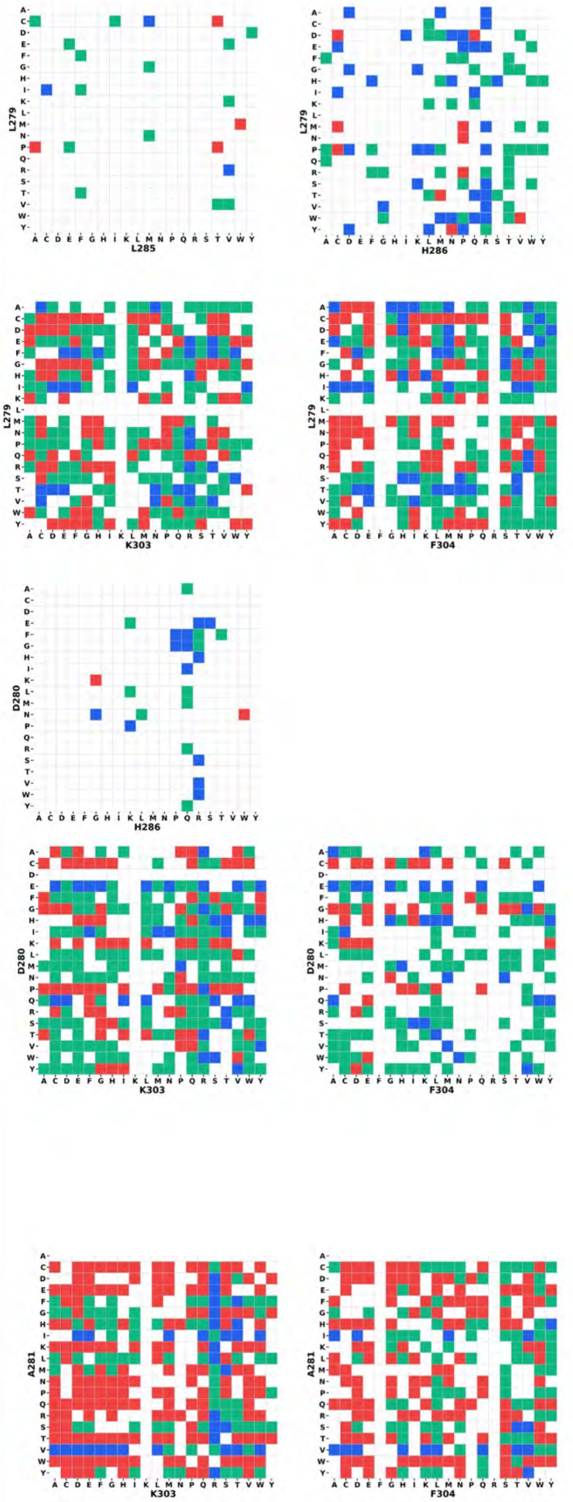

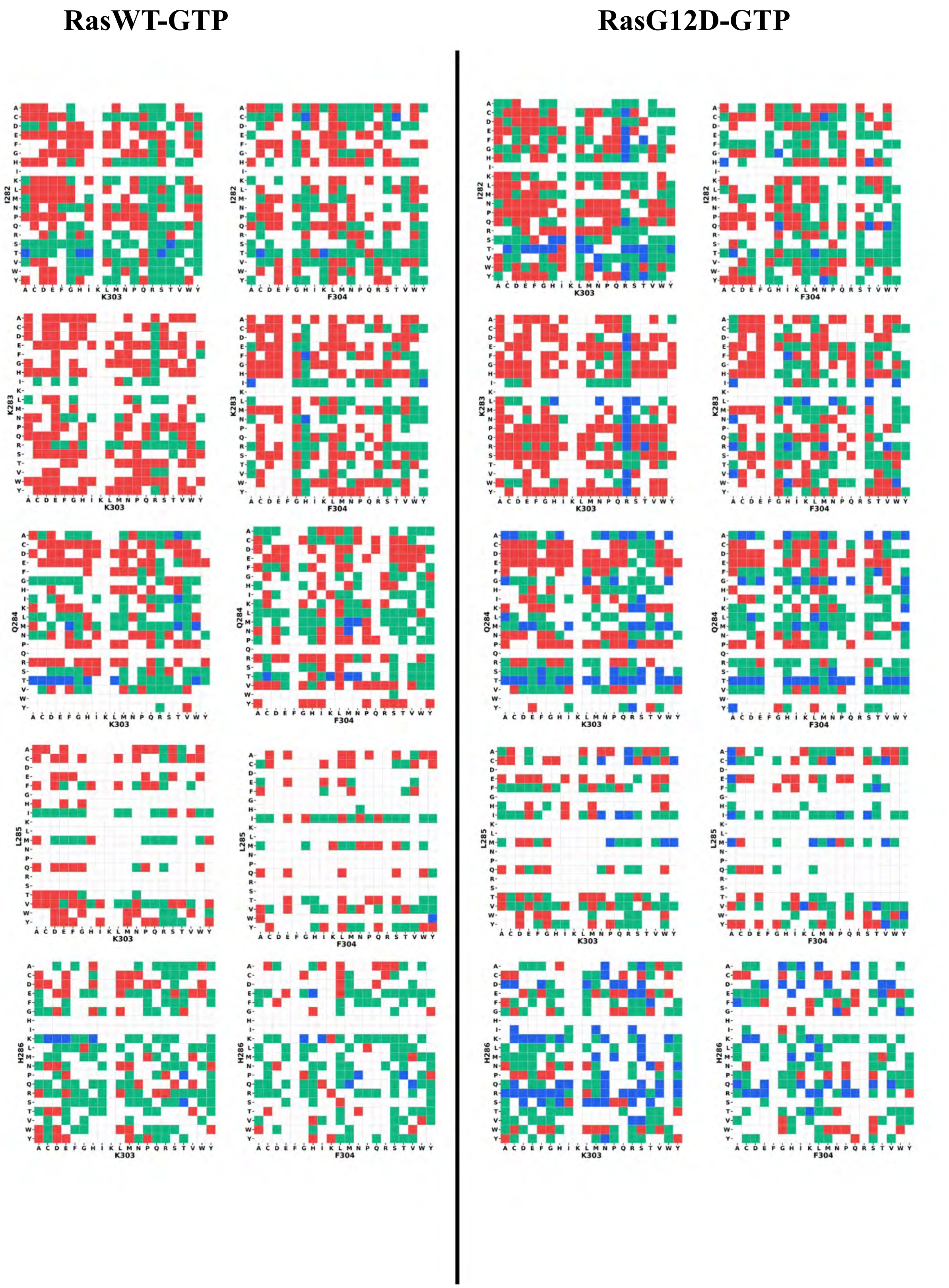


**Figure S8: Additivity/non-additivity of double mutations detected in the RasWT-GTP (left) and RasG12D-GTP (right) interactions.** The squares are color coded according to the mutation interaction energy $\Delta G_i$ calculated in equation 4. Additive mutations are shown in green, mutations with negative epistasis in blue, with positive epistasis in red.

#### Supplementary Tables and Legends

**Supplementary Table S1** $K_D$ and ΔΔGbind values for Nore1A variants interacting with RasWT-GTP and RasG12D-GTP

| Nore1A mutant | RasWT-GTP | | RasG12D-GTP | | Specificity shift* |
|---|---|---|---|---|---|
| | Kd, μM | $\Delta\Delta G_{bind}$, kcal/mol | Kd, μM | $\Delta\Delta G_{bind}$, kcal/mol | |
| WT | 0.22 ± 0.06 | 0 | 0.55 ± 0.22 | 0 | - |
| Q284T | 0.0017 ± 0.009 | -2.86 | 0.0040 ± 0.0016 | -2.91 | 1.06 |
| L277Y | 0.18 ± 0.031 | -0.11 | 0.27 ± 0.07 | -0.41 | 1.66 |
| L279V | 0.089 ± 0.040 | -0.54 | 0.219 ±0.074 | -0.55 | 1.06 |
| Q284M | 0.050 ± 0.023 | -0.89 | 0.084 ±0.025 | -1.11 | 1.48 |
| I282T | 0.097 ± 0.045 | -0.48 | 0.187 ± 0.043 | -0.64 | 1.30 |
| L277H | 0.186 ± 0.030 | -0.09 | 0.380 ± 0.079 | -0.22 | 1.08 |
| H286S | 0.294 ± 0.049 | 0.188 | 0.213 ± 0.086 | -0.56 | 3.55 |
| A281M | 0.396 ± 0.078 | 0.37 | 1.19 ± 0.44 | 0.45 | 1.14 |
| L221R | 1.12 ± 0.18 | 1.00 | 0.99 ± 0.44 | 0.34 | 3.06 |
| D280H | 0.415 ± 0.084 | 0.4 | 0.53 ± 0.14 | -0.022 | 2.04 |
| K283M | 1.97 ± 1.43 | 1.35 | 1.54 ± 0.82 | 0.60 | 3.9 |
| I282Q | 0.44 ± 0.08 | 0.43 | 1.11 ± 0.17 | 0.41 | 1.0 |
| I282C | 0.59 ± 0.24 | 0.61 | - | - | - |
| K303H | 2.65 ± 2.02 | 1.53 | 4.93 ± 2.35 | 1.28 | 1.5 |
| H286K | 0.033 ± 0.007 | -1.13 | 0.050 ±0.009 | -1.41 | 1.6 |
| L221A** | | 1.26 | | | |
| L221D** | | 1.87 | | | |
| F234A** | | 0.79 | | | |
| D280A** | | 0.30 | | | |
| K283A** | | 2.26 | | | |
| Q284A** | | 0.26 | | | |
| K303A** | | 1.55 | | | |

***Specificity shift was calculated as $K_D$(Nore1A mut to RasWT)/$K_D$(Nore1A WT to RasWT)· $K_D$(Nore1A WT to RasG12D)/$K_D$(Nore1A mut to RasG12D)**

****The data was taken from Stieglitz, B. et al (7)**

#### Multimedia File Legends (Videos / Datasets)

The videos and datasets could be downloaded from:

https://zenodo.org/records/23944358?token=eyJhbGciOiJIUzUxMiJ9.eyJpZCI6IjJhNzU0MDQ0LTVlNGMtNDFhMC05MzRkLTY5YjBkYWNhZmY4NiIsImRhdGEiOnt9LCJyYW5kb20iOiIzZWUzZmIwOWI2ODQ1ZWIxMmQ0YTk1ZWI5NGVkNGUzMyJ9.54pYcmvqTsiL0EVTwY6sWgpW6237BV53WPMB2THJs0pTHFwZB8bH6WdegtJQVd_zy5lO08FL2jQx_RrZuB4NXw

**Movie S1 (Nore1AWT_RasWT.avi):** Molecular dynamics trajectory showing the structural dynamics of wild-type Nore1A (grey cartoon) in complex with wild-type Ras (cyan cartoon). The bound $Mg^{2+}$ ion is depicted as a green sphere, and residue position 12 of Ras is highlighted as a red sphere.

**Movie S2 (Nore1AWT_RasG12D.avi):** Molecular dynamics trajectory showing the interface dynamics of wild-type Nore1A (grey cartoon) bound to mutant Ras(G12D) (cyan cartoon). The bound $Mg^{2+}$ ion is shown as a green sphere, and the G12D mutation site (position 12) is highlighted as a red sphere.

**Movie S3 (Nore1AH286K_RasWT.avi):** Molecular dynamics trajectory illustrating the interaction dynamics of the Nore1A(H286K) variant (grey cartoon) complexed with wild-type Ras (cyan cartoon). The bound $Mg^{2+}$ ion is shown as a green sphere, and residue position 12 of Ras is indicated by a red sphere.

**Movie S4 (Nore1AH286K_RasG12D.avi):** Molecular dynamics trajectory capturing the interface behavior of the Nore1A(H286K) variant (grey cartoon) bound to mutant Ras(G12D) (cyan cartoon). The bound $Mg^{2+}$ ion is shown as a green sphere, and the G12D mutation site (position 12) is indicated by a red sphere.

**Dataset 1. Normalisation dataset and leave-one-out predictions.**
The experimental dataset includes experimentally measured $\Delta\Delta G_{bind}$ values and normalized enrichment values for Nore1A variants interacting with RasWT-GTP and RasG12D-GTP. The enrichment values were obtained from four FACS gates. These data were used to train and validate an Extra Randomized Trees model. For each Nore1A variant, the corresponding $\Delta\Delta G_{bind}$ value predicted using leave-one-out cross-validation is provided.

**Dataset 2. Predicted $\Delta\Delta G_{bind}$ values for the 4 PPIs (Nore1A single and double mutants binding to RasWT and RasG12D in the GTP- and GDP-bound states).** The dataset includes all detected single and double mutants after filtering out mutations that were detected fewer than 10 times in the naïve library and mutations detected in less than 3 gates. For each mutant, the table lists the mutation name, the four normalized enrichment values (natural-log scale), the predicted $\Delta\Delta G_{bind}$ from the model trained on the normalized dataset, and the estimated prediction error ($\sigma$).

**Dataset 3. Raw enrichments values for single and double Nore1A mutants interacting with RasWT and RasG12D in the GTP- and GDP-bound states.** The dataset includes all detected single and double mutants before any filtering was applied. Raw enrichment values were calculated according to equation 2 in the manuscript.
